# A double-negative prostate cancer subtype is vulnerable to SWI/SNF-targeting degrader molecules

**DOI:** 10.1101/2024.03.24.586276

**Authors:** Phillip Thienger, Irene Paassen, Xiaosai Yao, Philip D. Rubin, Marika Lehner, Nicholas Lillis, Andrej Benjak, Sagar R. Shah, Alden King-Yung Leung, Simone de Brot, Alina Naveed, Bence Daniel, Minyi Shi, Julien Tremblay, Joanna Triscott, Giada Andrea Cassanmagnago, Marco Bolis, Lia Mela, Himisha Beltran, Yu Chen, Salvatore Piscuoglio, Haiyuan Yu, Charlotte K Y Ng, David A. Quigley, Robert L. Yauch, Mark A. Rubin

## Abstract

Proteolysis targeting chimera (PROTAC) therapies degrading SWI/SNF ATPases offer a novel approach to interfere with androgen receptor (AR) signaling in AR-dependent castration-resistant prostate cancer (CRPC-AR). To explore the utility of SWI/SNF therapy beyond AR-sensitive CRPC, we investigated SWI/SNF-targeting agents in AR-negative CRPC. SWI/SNF targeting PROTAC treatment of cell lines and organoid models reduced the viability of not only CRPC-AR but also WNT-signaling dependent AR- negative CRPC (CRPC-WNT). The CRPC-WNT subgroup represents 11% of around 400,000 cases of CRPC worldwide who die yearly of CRPC. We discovered that SWI/SNF ATPase SMARCA4 depletion interfered with the master transcriptional regulator TCF7L2 (TCF4) in CRPC-WNT. Functionally, TCF7L2 maintains proliferation via the MAPK signaling axis in this subtype of CRPC. These data suggest a mechanistic rationale for interventions that perturb the DNA binding of the pro-proliferative TCF7L2 transcription factor (TF) and/or direct MAPK signaling inhibition in the CRPC-WNT subclass of advanced prostate cancer.

**Statement of significance:** Androgen receptor (AR)-negative prostate cancer (PCa) remains a clinical challenge due to the lack of targeted therapeutic options. Here, we identified a lineage-defining molecular axis in a subtype of AR- negative PCa, accounting for around 10% of castration-resistant PCa (CRPC) that can be interfered with by SWI/SNF-targeting agents.

## Introduction

Treatment-induced shifts in cancer cell identity, known as lineage plasticity (LP), lead to the emergence of tumors that may have little to no resemblance to the treatment-naïve tumors. The increased and earlier use of potent targeted cancer therapies are responsible for the emergence of aggressive, “plastic,” and untreatable cancers. In prostate cancer (PCa), LP can manifest when androgen receptor (AR)-driven adenocarcinoma (CRPC-AR) differentiates into AR-negative CRPC, which lacks both canonical AR signaling and neuroendocrine differentiation markers (double-negative CRPC, aka DNPC) or acquires neuroendocrine features (CRPC-NE)^1–3^.

The epigenetic chromatin remodeling machinery complex switch/sucrose non-fermentable (SWI/SNF) orchestrates pluripotency and differentiation in embryonic stem cells^4^, indicating its potential to maintain self-renewal in cancer and modulate lineage plasticity. In line with this, the SWI/SNF complex is mutated in over 20% of cancers^5^. However, in PCa, genomic alterations in the SWI/SNF complex are rare. Regardless, we and others have observed the dysregulation in SWI/SNF ATP-dependent helicases SMARCA2 (BRM) and SMARCA4 (BRG1) expression levels in CRPC^6,7^. In non-small lung cancer (NSCLC), alterations in SWI/SNF ATPase expression (mainly loss of SMARCA4) have led to the discovery of a synthetic lethal relationship^8–10^. In CRPC-AR, degradation of the SWI/SNF catalytic ATPase subunits (SMARCA2, SMARCA4) compacts cis-regulatory elements bound by AR-associated transcription factors (TFs) leading to drastic decrease of PCa proliferation^11^.

Another critical pathway in cancer, especially during developmental processes, is the wingless and int-1 (WNT) pathway, which is partially regulated by the SWI/SNF complex^12–14^. Aberrations in WNT signaling are especially prominent in colorectal cancer but also emerge in other cancer types, such as CRPC^15–17^. This signaling pathway can be divided into the canonical (β-Catenin-dependent) and non-canonical (β- Catenin-independent) axis^18^. In PCa, genetic changes in canonical WNT pathway genes are found in up to 22% of CRPC cases, while also non-canonical WNT signaling is altered in advanced PCa^19,20^. Further, the PCa stroma is known to secrete WNT proteins that activate WNT signaling in tumor cells to promote therapy resistance^21,22^. Recently, Tang et al. identified a subclass of CRPC, termed CRPC-WNT, that has traits of double-negative PCa (DNPC) but is enriched for mutations in WNT signaling pathway genes, accompanied by strong pathway activation through the TF TCF7L2, among others^23^.

In this study, we report that targeting SMARCA2/4 downregulates this lineage-defining WNT signaling signature^23^ in CRPC-WNT patient-derived organoids. Indeed, we found that SMARCA4 depletion led to the chromatin closure at TCF7L2 DNA binding motifs and the downregulation of TCF7L2 itself in CRPC-WNT. Further, we provide evidence that this downregulation of TCF7L2 is facilitated through closure of an active intragenic enhancer. By performing chromatin immunoprecipitation sequencing, we narrowed down the function of TCF7L2 to be involved in pro-proliferative pathways such as RAS and MEK signaling by binding to relevant gene promotors. We functionally validated that CRPC-WNT are addicted to these signaling pathways and that SMARCA2/4 degradation by A947 treatment reduces the protein levels of known MEK downstream targets.

This is in line with findings that described MEK signaling as a dependency in DNPC^1^. In summary, we found TCF7L2 to be a primary driver of CRPC-WNT, which is positively regulated by SMARCA4- dependent SWI/SNF activity to drive proliferative pathways. This study strengthens the evidence that the SWI/SNF complex plays a crucial role in advanced PCa and can be therapeutically exploited beyond AR- driven PCa adenocarcinoma.

## Results

### SMARCA2/4 is a vulnerability in DNPC patient-derived organoids (PDO)

In prior work, we discovered overexpression of SMARCA4 correlates with PCa progression, particularly neuroendocrine prostate cancer^6^. The testing of SMARCA2/4 targeting agents in PCa had been restricted to standard cell lines, covering only a limited representation of the commonly encountered genomic landscape and progression states seen in patients with CRPC. We posited that AR-negative CRPC may also manifest sensitivity to SWI/SNF ATPase inhibition. To address this, we used a novel PROTAC degrader, A947, that co-binds the SMARCA2/SMARCA4/PBRM1 bromodomains and the von Hippel-Lindau (VHL) ubiquitin ligase^9^ (**Fig.1a**). This molecule has slight selectivity for SMARCA2 degradation, however at the concentrations used here it is equally potent for degradation of both ATPases within 1 hour in HEK293 cells (**Fig.1b**).

A947 was tested on a panel of PCa models, including established and organoid-derived cell lines and the non-neoplastic prostate line RWPE-1 (**Fig. 1c, Supplementary Figure 1a** and **b**). The four CRPC subclasses described by Tang et al.^23^, AR-dependent (CRPC-AR) (n=6), “WNT-driven” (CRPC-WNT) (n=4), neuroendocrine prostate cancer (NEPC) (n=4), and “stem-cell-like” (CRPC-SCL) (n=7) were treated for seven days with A947 in a dose response (**Supplementary Figure 1a**). We confirmed that AR-dependent cell models are particularly sensitive to SWI/SNF degradation^11^. In addition, we discovered that CRPC-WNT models were fully or partially responding to A947 (area under the curve (AUC<250)) (**Fig.1c, Supplementary Figure 1a**). All PCa model systems tested were non-responsive to negative control A858 epimer (SMARCA-binding control) as expected **(Supplementary Figure 1a**). A947 was able to degrade all predicted targets, SMARCA2, SMARCA4 and PBRM1 in all four CRPC-WNT models (**Supplementary Figure 1c**). Similar anti-proliferative responses as to A947 were observed with SMARCA2/4 PROTAC AU-15330 and inhibitors FHD-286 and BRM014 in all four CRPC-WNT and CRPC-AR but not in other subtypes (**Supplementary Figure 2a, b and c**). Next, we checked for SMARCA2, SMARCA4 and lineage-defining marker expression in selected organoids, as well as cell lines and found the expected marker gene expression pattern, while the expression for SMARCA4 was overall higher than the expression of SMARCA2 in most models, also the ones that showed a response to A947-treatment (**Fig. 1d**).

**Fig. 1.**
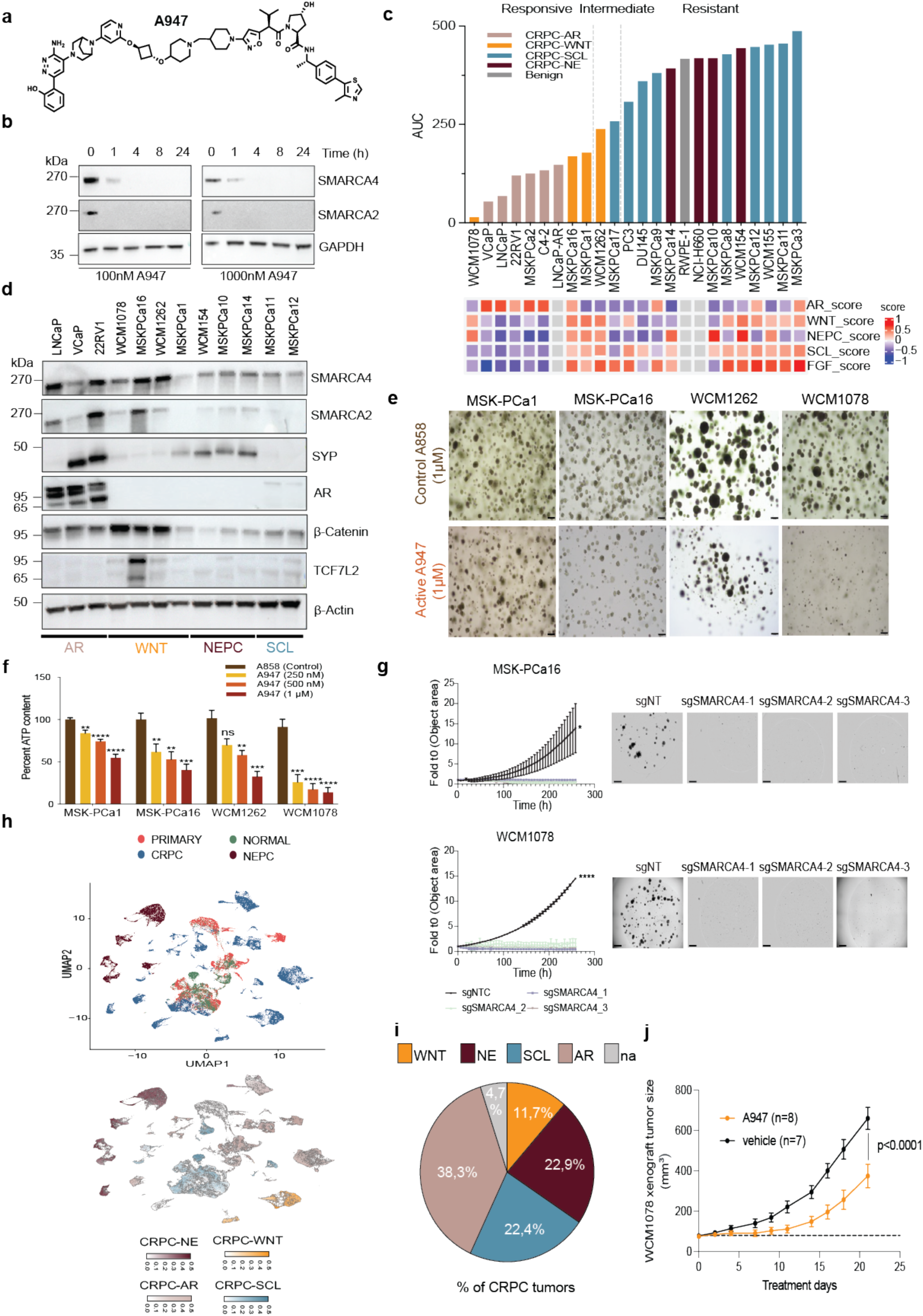
Drug screen identifies SMARCA2/4 as a vulnerability in DNPC cell models. a, Structure of A947 SMARCA2/4 PROTAC degrader. b, Immunoblot of indicated proteins in HEK293 cells treated with A947 (0.1µM or 1µM) or DMSO over indicated time course. GAPDH represents loading control and is probed in a representative immunoblot (*n*=2 independent immunoblots). c, Area under the curve (AUC) of A947 dose response curves (see Supp. figure 1a and b) in a panel of human-derived prostate cancer or normal cell lines after 7 days of treatment. Viability was assessed using Celltiter Glo 2.0. Heatmap indicates gene-set scores per cell model using Tang et al.^23^ scores. Data is representative of at least *n* = 3 independent experiments. d, Immunoblot of indicated proteins in a panel of PCa organoids representing different phenotypes. β- Actin serves as a loading control and is probed in a representative immunoblot (*n*=1). e, Brightfield microscopy of indicated PCa organoids after 10 days of treatment with A947 (1µM) or epimer control A858 (1µM). Scale bars: 20 μm. f, Proliferation of indicated PCa organoids after 10 days of treatment with A947 (at 0.25µM, 0.5µM, 1µM) or epimer control A858 (1µM) measured by Celltiter Glo 3D (n = 3 independent biological experiments). Data are presented as mean values +/− SEM and analyzed using unpaired Students t-test (*p < 0.05, **p < 0.01, ***p < 0.001, ****p < 0.0001). Data is representative of at least *n* = 3 independent experiments. g, Spheroid formation of indicated PCa organoids transduced with CRISPRi-Cas9 guide RNA (sgRNA) against SMARCA4 measured by live imaging using Incucyte SX5. N = 3 independent experiments. Brightfield microscopy of indicated PCa organoids at Incucyte assay endpoint, Scale bars: 800 μm. Data are presented as mean values +/− SEM and analyzed using two-way ANOVA (*p < 0.05, **p < 0.01, ***p < 0.001, ****p < 0.0001). Data is representative of *n* = 2 independent experiments. h, UMAP plot showing disease classification (left) and relative expression of indicated signature gene scores (right) in 74 samples from six distinct PCa scRNAseq studies^2,25–29^. Side annotations indicate the AR score, NE score, stem cell-like (SCL) score, and WNT score, as determined by Tang et al.^23^, compared with pathology classification and molecular subtypes of each sample. False discovery rate (FDR) <0.05. i, Percentage of indicated signatures found among all CRPC classified samples displayed as a pie chart. Not announced (na). j, Tumor growth of WCM1078 PDX subline treated with vehicle (n=7) or 40mg/kg A947 (n=8) in xx mice. Data are presented as mean values +/− SEM and analyzed using two-way ANOVA (p < 0.0001).

Organoid formation was drastically reduced in CRPC-WNT organoids after 7 days of A947 treatment (**Fig. 1e and f; Supplementary Figure 2d**). Assessment of growth kinetics using live-cell imaging in CRPC-WNT 2D lines MSK-PCa16 and WCM1078 showed significant reduction in cell confluence over time to a single dose (1µM) of A947 compared with control epimer A858 (**Supplementary Figure 2e**). Strikingly, competition of A947 with a free VHL ligand (VL285) rescued growth inhibitory effect dose dependently (**Supplementary Figure 2f**). Notably, levels of cleaved PARP1 (Asp214) were increased in the CRPC-WNT lines, WCM1078 and MSK-PCa16 upon treatment with 1µM A947 indicating activation of the intrinsic pathway of cell death (**Supplementary Figure 2g**).

To estimate the necessity of SMARCA2/4 activity in the CRPC-WNT, we performed individual siRNA- mediated knockdown of these two main A947 targets in CRPC-WNT models, a CRPC-NE model and CRPC-SCL model. All CRPC-WNT models showed a strong growth inhibitory effect upon SMARCA4 knockdown but only minimally responded to the knockdown of SMARCA2. CRPC-NE model WCM154 and CRPC-SCL model WCM155, however, were unaffected by either siRNA knockdown (**Supplementary Figure 3a, b)**. SMARCA4 inhibition has been reported to be synthetic lethal with PTEN loss in PCa^24^. However, only two of the four CRPC-WNT models harbor PTEN deletions, indicating that the response to SMARCA4 degradation may be independent of its PTEN status^23^. To rule out siRNA- mediated off-target effects, we elucidated the isolated effects of SMARCA4 depletion on cell growth using CRISPR-Cas9 sgSMARCA4 transduced organoids. We confirmed similar growth inhibition and cell-killing effect to A947 treatment upon SMARCA4 knockdown using CRISPRi in all four CRPC-WNT lines (**Fig. 1g; Supplementary Figure 3c, d**). In conclusion, we identified an AR-negative subtype of CRPC that is dependent on the SWI/SNF ATPase SMARCA4 in vitro.

### CRPC-WNT is a clinically relevant subset of advanced CRPC

To determine how frequently the CRPC-WNT phenotype identified by Tang et al.^23^ is seen clinically, we used publicly available single-cell sequencing (scRNA-seq) data from PCa cohorts (which include normal prostate/primary PCa (n=45), CRPC/NEPC (n=29) samples in total^2,25–29^). We found that the four signatures described by Tang et al.^23^ could be identified in distinct clusters, especially the CRPC-WNT signature, which appeared in two distinct subclusters of CRPC separate from all other clusters (**Fig. 1h**). All remaining signatures, CRPC-AR, CRPC-NE, and CRPC-SCL, were more broadly distributed. Surprisingly, the CRPC-SCL signature was not only highly expressed in CRPC cases but also in normal and primary PCa (**Fig. 1h**). Overall, the signatures for CRPC-AR, CRPC-WNT, CRPC-NE, and CRPC- SCL account for 38.3%, 11,7%, 22,9% and 22.4% of all CRPC cases, respectively. This underpins the clinical relevance of these signatures. These signatures could not characterize 4.7% of CRPC cases, indicating that additional rare phenotypes of CRPC may exist (**Fig. 1i**). This highlights that CRPC-WNT is a highly clinically relevant subtype of PCa for which there is no viable treatment option available. All these findings encouraged us to investigate the effect of SMARCA2/4 degradation on CRPC-WNT in vivo

### Treatment with SMARCA2/4 PROTAC leads to CRPC-WNT tumor growth delay in vivo

To assess the effect of SMARCA2/4 degradation in vivo, we generated a mouse-adapted patient-derived xenograft subline from the WCM1078 CRPC-WNT model. Since WCM1078 did not reliably grow in vivo, we had to create a subline from a single WCM1078 tumor that grew only in 1 out of 5 injected mice within 6 months. Tumor cells were expanded in vitro and injected again into mice until visible tumors formed. From these tumors, we generated cryobits. These cryobits were subcutaneously transplanted into 20 mice. 16/20 of the animals developed tumors; 15 were taken into the study, and the others were excluded. A single dose of A947 treatment (40mg/kg) or vehicle was given to 8 animals or 7 animals, respectively, when tumors reached 60-80mm^3^ (**Supplementary Figure 4a**). Before treatment, we checked the potential of A947-treatment to reduce mouse SMARCA4 paralog in vitro. We found that the compound is active in murine lung adenoma LA4 cells (**Supplementary Figure 4b**). We observed a significant growth delay in the A947-treatment condition (n=8) compared to vehicle control (n=7), which aligns with the findings by Xiao et al.^11^ in CRPC-AR (**Fig. 1j, Supplementary Figure 4c and d**). Moreover, the mice showed no signs of aberrant behavior or reduction in body mass throughout treatment (**Supplementary Figure 4e**). Histopathological assessment indicated that all examined tissues were unremarkable (**Supplementary Figure 4f**), except for histopathologic findings in the kidney. Chronic renal interstitial fibrosis and tubular atrophy were identified in two A947 treated mice where this change was mild (C14) to moderate (C11) (**Supplementary Figure 4g**). As this change is also known to occur spontaneously in laboratory mice (chronic nephropathy), it remains unknown if this lesion is related to the A947 treatment. Next, we assessed the levels of SMARCA4 in the tumors harvested at the endpoint. Quantifying the western blot signal showed a significant average reduction in SMARCA4 protein in the A947-treated tumors compared with control despite the single-time treatment (**Supplementary Figure 4h**). This indicates that A947 is highly active in in vivo and can reduce CRPC-WNT tumor growth. In summary, we found that CRPC-WNT is highly responsive to a single treatment of A947 throughout a 21-day tumor growth period in vivo. These findings led us to investigate the underlying molecular mechanisms regulated by SMARCA4 in CRPC-WNT, which accounts for 11.7% of all CRPC patients.

### A lineage-defining WNT program is mitigated by SMARCA2/4 degradation in CRPC-WNT

To untangle the transcriptomic and heterogeneous changes upon treatment with A947 in CRPC-WNT, we utilized single-cell RNA sequencing (scRNAseq) by SORT-seq^30^. WCM1078 and MSK-PCa16 were treated for 72h with 1µM A947 or control epimer A858. A total of 1133 WCM1078 cells and 1183 MSK- PCa16 cells in the A858-treated condition and a total of 1184 WCM1078 cells and 997 MSK-PCa16 cells in the A947-treated condition passed quality control. Both PDOs demonstrated homogenous profiles with separate clusters forming based on treatment (**Fig. 2a and e**). As expected, CRPC-WNT signature score^23^ was homogenously expressed in the A858 conditions, while this signal was significantly reduced after A947 treatment (**Fig. 2b, c, f and g**). High expression in A858 treated cells was observed for an additional WNT signature score based on colorectal cancer (CRC_WNT_score; compiled of the genes *"LGR5", "AXIN2", "ASCL2", "OLFM4", "SLC12A2", "GKN33P", "NKD1", "WIF1"*) although the signal was more heterogeneously expressed (**Supplementary Figure 5a and c**). This CRC-WNT score signal was significantly reduced by A947-treatment, in line with the reduced CRPC- WNT signature (**Supplementary Figure 5b and d**). However, besides these two specific WNT signatures we did not observe significant changes in other WNT-pathway signatures besides one LEF- related signature (**Supplementary Figure 5i and j**). This indicates that the genes that define those signatures might not be tied to canonical WNT signaling in CRPC-WNT. Therefore, we posit that the CRPC-WNT score is a refined lineage-specific signature comprised of known WNT TFs, such as TCF7, TCF7L2, and LEF1, that might have non-canonical functions.

**Fig. 2.**
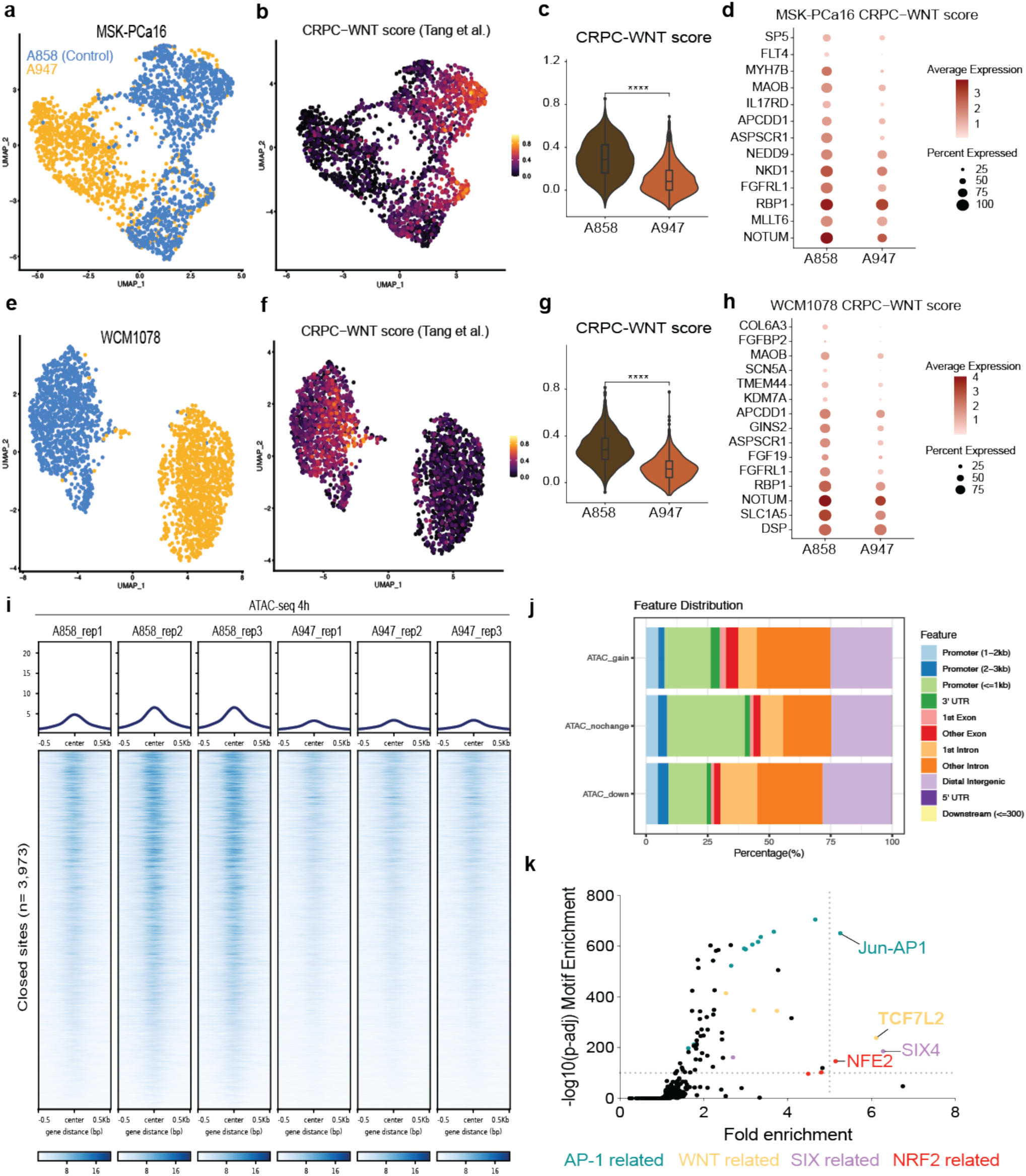
SMARCA2/4 degradation leads to strong downregulation and chromatin compaction of CRPC-WNT lineage-characterizing genes. a, UMAP plot of MSK-PCa16 organoids treated with either 1µM A858 (blue) or 1µM A947 (yellow) for 72h. b, UMAP plot of MSK-PCa16 organoids treated with either 1µM A858 or 1µM A947 for 72h displaying CRPC-WNT signature score^23^. c, Violin plot of MSK-PCa16 organoids treated with 1µM A858 or 1µM A947 for 72h displaying CRPC- WNT signature^23^ signature score. Analyzed using the Wilcoxon test (****p < 0.0001). d, Bubble plot indicative of expression levels of top 10 deregulated CRPC-WNT signature genes^23^ in MSK-PCa16 organoids treated with 1µM A858 or 1µM A947 for 72h. e, UMAP plot of WCM1078 organoids treated with 1µM A858 (blue) or 1µM A947 (yellow) for 72h. f, UMAP plot of WCM1078 organoids treated with 1µM A858 or 1µM A947 for 72h displaying CRPC-WNT signature score^23^. g, Violin plot of WCM1078 organoids treated with 1µM A858 or 1µM A947 for 72h displaying CRPC-WNT signature^23^ signature score. Analyzed using the Wilcoxon test (****p < 0.0001). h, Bubble plot indicative of expression levels of top 10 deregulated CRPC-WNT signature genes^23^ in WCM1078 organoids treated with 1µM A858 or 1µM A947 for 72h. i, ATAC-seq read-density tornado plots from WCM1078 organoids treated with 1µM A858 or 1µM A947 for 4h (*n* = 3 biological replicates). Venn diagram indicating lost regions. j, Genome-wide changes in chromatin accessibility upon A947-treatment for 4 h in WCM1078 organoids along with genomic annotation of sites that gain (gained) or lose accessibility (lost) or remain unaltered (unchanged). k, Motifs enriched in depleted peaks from WCM1078 treated for 4h with 1µM A947 identified using HOMER on ATAC-seq.

Notably, other CRPC subtype signature scores (CRPC-AR, CRPC-SCL, CRPC-NE) were barely expressed in the CRPC-WNT organoids (**Supplementary Figure 5e** and **f)**. Notably, The CRPC-SCL and CRPC-NE scores were also downregulated by A947 treatment compared with A858 treatment, but the positive clusters had low base levels compared with the CRPC-WNT positive cluster (**Supplementary Figure 5e** and **f**). This was confirmed by bulk RNA-seq data on WCM1078 after 24h and 48h with A947 (**Supplementary Figure 6a**). Overall, several individual genes that define CRPC-WNT, as characterized by Tang et al.^23^, were downregulated upon treatment with A947 in both organoid models (**Fig. 2d and h**). SCENIC pathway analysis identified decreased activity of multiple TFs upon A947-treatment^31^. In both, WCM1078 and MSK-PCa16, activity of several TFs has been reduced; namely activity of FOX, TCF/WNT, ETV and AP-1 family members showed decreased activity (**Supplementary Figure 5g** and **h)**. In agreement, bulkRNA-seq data of WCM1078 after A947 treatment showed decreased expression of a subset of the Top 25 highest-ranked transcription factors in CRPC-WNT^23^, among the most downregulated TFs: KLF2, TCF7L1, TCF7L2, SOX13, SOX4, RUNX3 and LEF1 (**Supplementary Figure 6b**). Gene ontology analysis of oncogene C6 signature revealed a downregulation of MAPK signaling signatures terms (ERBB2_UP.V1_UP, MEK_UP.V1_UP) upon A947 treatment in WCM1078 cells (**Supplementary Figure 5i**). Moreover, we found that in MSK-PCa16 scRNA-seq the most downregulated C6 oncogene pathways were associated with WNT (LEF1_UP.V1_UP) and MAPK signatures (KRAS.600_UP.V1.UP, KRAS.KIDNEY_UP.V1_UP) (**Supplementary Figure 5j**).

Gene set enrichment analysis of bulk RNA-seq data of WCM1078 after 48h treatment revealed that multiple oncogene C6 signatures, associated with mitogen-activated protein kinase (MAPK)-KRAS-MEK signaling, were downregulated by A947-treatment (**Supplementary Figure 6c**). This data validates our previous finding in the scRNA-seq analysis and points to a potential MAPK-associated proliferative axis in CRPC-WNT which is disrupted by A947 treatment. This might be a downstream effect of the loss of the lineage-defining CRPC-WNT signature. To gain further insight into the epigenetic orchestration of this process, we examined chromatin accessibility in CRPC-WNT upon treatment with A947.

### SMARCA2/4 degradation leads to closure of TCF/LEF chromatin binding sites in CRPC-WNT

The SWI/SNF complex mediates nucleosomal DNA packaging and is actively involved in regulating gene expression of multiple programs that can be crucial for cell survival. To mechanistically exploit changes in chromatin accessibility, we profiled the changes mediated by A947 treatment using the assay for transposase-accessible chromatin followed by sequencing (ATAC-seq). Within 4 hours, we found a near- complete loss at 3,979 sites in WCM1078 (CRPC-WNT), while only 80 sites were gained compared with A858 treatment (**Fig. 2i**). The compaction of lost sites in WCM1078 organoids was comparable to what has been found in CRPC-AR cell lines upon the treatment with SMARCA2/4 PROTAC^20^ (**Supplementary Figure 6d**). Overall, we see that over 50% of A947-treatment compacted sites are associated with intronic and distal intergenic regions like what has been observed in CRPC-AR^20^ (**Fig. 2j**). Transcription factor motif analysis of A947 lost sites revealed lost motif accessibility of several CRPC-WNT driving TFs, among others: Jun-AP1, TCF, LEF, SIX, NFE, FOX, and SOX motifs (**Fig. 2k, Supplementary Figure 7a**). The top depleted motifs upon A947 treatment are TCF7L2-related, which has been identified as the most active TF in CRPC-WNT^23^. Also, the motif Jun-AP1 is associated with WNT signaling factors since c-Jun is known to form a pro-proliferative complex with β-Catenin and TCF TFs, including TCF7L2^32–34^.

Moreover, SIX TFs are known to directly interact with TCF7L2 and drive PCa cell plasticity via WNT signaling^35–37^. The fact that we see FOX motifs downregulated aligns with the findings made in CRPC-AR, indicating also the on-target selectivity of A947^20^. When performing GSEA of the ATAC-seq and ATAC-seq/RNA-seq overlap results, we found that gene sets associated with MEK signaling were found to be downregulated by A947-treatment (**Supplementary Figure 7b-e**). These findings indicate that TCF/LEF signaling motifs are direct targets of the SWI/SNF complex remodeling, potentially regulating several downstream effectors associated with carcinogenesis and proliferation.

### TCF7L2 is a dependency in CRPC-WNT

To test if WNT signaling and TCF7L2 expression is critical for CRPC-WNT survival we performed siRNA-mediated knockdown experiments. MSK-PCa16, which has an amplification in TCF7L2, indeed showed a growth inhibitory effect upon siTCF7L2 transfection (**Fig. 3a**). This phenotype was rescued by the transfection of a transient TCF7L2 rescue construct indicating the effect of siRNA on cell confluence is not due to off-target effects (**Fig. 3a, Supplementary Figure 8a**). In line with this, spheroid formation of other CRPC-WNT models was reduced when depleting TCF7L2 and β-Catenin (CTNNB1) using siRNA (**Fig. 3b, Supplementary Figure 8b**). Notably, CRPC-NE and CRPC-SCL models which express TCF7L2 were unaffected by siTCF7L2 (**Supplementary Figure 8c**). To assess potential bias from the organoid growth media, which, contains R-spondin 1 (RSPO1), an activator of WNT-signaling, we measured the growth of WCM1078 and MSK-PCa16 in the absence of RSPO1 upon knockdown of β- Catenin and TCF7L2. The growth phenotype with and without RSPO1 showed no significant differences when transfected with siRNA against WNT TFs, and the absence of RSPO1 only marginally affected general growth rates of these PDOs (**Supplementary Figure 8d**). Indeed, a recent study has identified that RSPO1 is not the inducing factor responsible for TCF7L2 expression in WNT-driven lineage progression^38^. This indicates that the WNT factors TCF7L2 and β-Catenin are potentially activated via non-canonical mechanisms or are constitutively active in CRPC-WNT indicated by harbored mutations in several WNT signaling factors^23^.

**Fig. 3.**
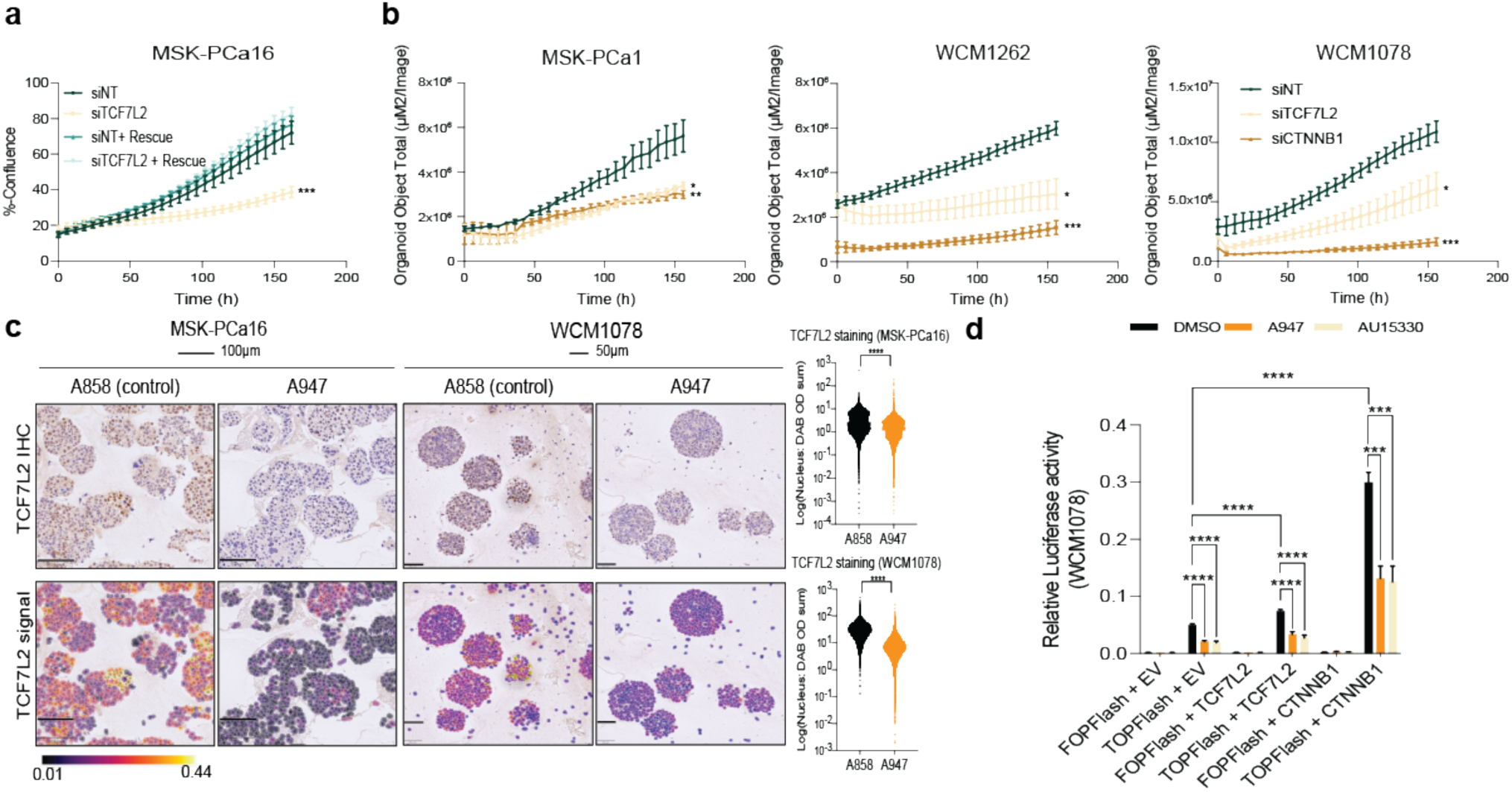
TCF7L2 is a dependency in CRPC-WNT. a, Growth data measured by live-cell imaging (Incucyte S3) upon transfection of indicated siRNA or plasmid (EV or rescue). Immunoblot of indicated proteins at 72h after transfection. GAPDH serves as loading control. Data are presented as mean values +/− SEM and analyzed using two-way ANOVA (*p < 0.05, **p < 0.01, ***p < 0.001). Data is representative of *n* = 2 independent experiments. EV: empty vector. b, Spheroid formation measured by live-cell imaging (Incucyte SX5) upon transfection of indicated siRNA. Data are presented as mean values +/− SEM and analyzed using two-way Anova (*p < 0.05, **p < 0.01, ***p < 0.001). Data is representative of *n* = 2 independent experiments. EV: empty vector. c, Immunohistochemistry (IHC) and staining intensity of TCF7L2 on indicated organoids upon treatment with 1µM A858 or 1µM A947 for 24h. Violin plot from TCF7L2 staining intensity analyzed using two-way ANOVA (*p < 0.05, **p < 0.01, ***p < 0.001, ****p < 0.0001). Scale: 100µm (MSK-PCa16), 50µm (WCM1078). d, TOPFlash TCF/LEF reporter assay measured after 48h upon treatment with indicated drugs in WCM1078 sublines. FOPFlash serves as a negative reporter control. Data are presented as mean values +/− SEM after normalization to internal Renilla control and analyzed using paired Students t-test (***p < 0.001, ****p < 0.0001). Data is representative of *n* = 2 independent experiments. EV: empty vector, OE: overexpression.

We next asked if SMARCA2/4 degradation has direct impact on the expression on WNT signaling TFs. Immunohistochemical staining of CRPC-WNT organoids treated with 1µM of control A858 or A947 for 24h revealed a strong downregulation of TCF7L2 signal (**Fig. 3c**). We found that also the SMARCA2/4 PROTAC AU-15330 reduced levels of TCF7L2 over time (**Supplementary Figure 8e**). Further, protein expression of TCF7L2, TCF7 and LEF1 was reduced over time upon A947-treatment, whereas β-Catenin levels were unaffected (**Supplementary Figure 8f**). This led us to assess the basis of TCF7L2 downregulation upon A947-treatment by looking at changes of chromatin structure.

### TCF7L2 expression is regulated through an active intragenic enhancer

As stated previously TCF7L2 binding sites are closing upon treatment with A947 and more strikingly, we observed downregulation on the protein level by immunohistochemistry and immunoblot in MSK-PCa16 and WCM1078 cells within 24h of treatment (**Fig. 3c, Supplementary Figure 8f**). Surprisingly, A947- treatment did not compact the TCF7L2 promoter loci, indicating closure of other regulatory sites lead to the downregulation of TCF7L2 levels. One of the earliest events in gene transcription is the activation of distal cis-regulatory enhancer regions and its associated transcription of eRNA. Since changes in chromatin structure are extremely rapid upon impairment of the SWI/SNF complex, it is not surprising to see the loss of accessibility of distal regulatory regions upon SWI/SNF complex inactivation^39,40^. Thus, we hypothesized that enhancer regions of TCF7L2 could be affected by A947-treatment.

To test our hypothesis, we explored the impact of A947-treatment on eRNA expression using the nuclear run-on followed by cap-selection assay (PRO-cap), which is the most sensitive method to identify active enhancers by measurement of endogenous eRNA transcription levels genome-wide at base-pair resolution^41^. Active enhancers loci can be precisely delineated by detecting active transcription start sites that are dependent on the associated core promoter sequences^42^ (**Supplementary Figure 9a**). When treating WCM1078 organoids for 24h with A858 or A947 (1µM) followed by PRO-cap, we found 1069 of eRNAs aka distal peaks (+/- 1kb) to be downregulated (blue) while only 2 distal peaks were upregulated (red) by A947-treatment (**Supplementary Figure 9b**). Next, we used the genomic search engine GIGGLE to identify and rank A947-treatment lost genomic loci shared between publicly available genome interval files^43^. These loci significantly overlapped with genomic sites bound by transcription factors associated with AP-1 (JUND, FOS, FOSL2, JUN), as well as FOXA1, ETV5, and TCF7L2 among the top 20 repressed distal peaks (**Supplementary Figure 9c**).

To identify potential promoter-enhancer loops regulating TCF7L2 expression, we exploited publicly available Hi-C data of 80 CRPC biopsy samples^44^. From these 80 samples, one patient with the highest CRPC-WNT gene expression score was selected for further inspection. Analysis of the TCF7L2 locus showed high contact frequency of the TCF7L2 promoter (chr10:112949674-112950536) with two intragenic regions (chr10:113087027-113087928 and chr10:113093268-113094092) that displayed high PRO-cap signal in WCM1078 treated with control epimer A858 (**Supplementary Figure 9d**). PRO-cap signal in these two intragenic regions after A947 treatment was significantly reduced. Decrease in PRO- cap signal was accompanied with decreased ATAC-signal and TCF7L2 binding to these intergenic regions. Intragenic regions were previously described as potential enhancer regions for TCF7L2^45,46^. Based on these findings we hypothesize that TCF7L2 regulates its own expression in CRPC-WNT patient by binding of an upstream intergenic enhancer region. Moreover, upon SMARCA2/4 degradation with A947 these potential enhancer loci reduce signal in PRO-cap, TCF7L2 ChIP-seq and ATAC-seq assays, indicating closure of those sites (**Supplementary Figure 9d**). This finding indicates that TCF7L2 expression is regulated by the SWI/SNF complex via maintenance of an intronic regulatory enhancer region in CRPC-WNT, similar to what has been reported for AR and FOXA1 in CRPC-AR^11^.

### TCF7L2 is not maintaining CRPC-WNT proliferation via traditional WNT signaling cues

For this we tested if A947-treatment interfered with transactivation of the TCF/LEF reporter TOPFlash^47^. We generated stable organoid lines from WCM1078 that express the multimerized TCF-binding site TOPFlash reporter or the negative control containing mutated TCF-binding binding sites (FOPFlash). To know if TCF7L2 or β-Catenin overexpression (OE) could rescue the expected downregulation of reporter signal by SMARCA2/4 PROTAC treatment, we overexpressed those two factors in the TOP/FOPFlash reporter organoids.

As expected, we found that both PROTACs, A947 and AU-15330, represses the TOPFlash reporter signal after 48h of treatment. While β-Catenin and TCF7L2 OE increased the reporter signal in DMSO control condition the PROTAC-treatment induced signal reduction could not be rescued in WCM1078 organoid lines (**Fig. 3d, Supplementary Figure 8g**). As we saw that A947-treatment represses the expression of multiple TCF and LEF TFs within 24h it is not surprising that OE of a single factor is not enough to restore the TCF/LEF reporter signal, when multiple TFs remain depleted (**Supplementary Figure 8f**).

To test whether CRPC-WNT are dependent on canonical WNT signaling, we treated CRPC-WNT organoids with three WNT inhibitors that have different modes of action (LGK974 (Porcupine inhibitor)^48^, iCRT14 (β-Catenin inhibitor)^49^, MSAB (β-Catenin inhibitor leading to its degradation)^50^. In addition, we treated the CRPC-NE model WCM154 and CRPC-AR model LNCaP, which should be “WNT- independent”, with these drugs. To our surprise, we found that all cell models used, including the “WNT- dependent” ones, did not respond to LGK974 or iCRT14. MSAB treatment led to decreased proliferation in all cell models tested, also the “WNT-independent” ones, at approximately the same concentration, indicting potential off-target effects of this drug (**Supplementary Figure 8h**). To see if pathway activation would have a stronger effect, we tested the WNT signaling agonist CHIR99021 (GSK3β inhibitor) in the CRPC-WNT model. Despite a slight increase in proliferation up to 1µM in MSK-PCa16 and WCM1262 the CRPC-WNT models were unresponsive to this agonist like we have observed with WNT agonist RSPO1 (**Supplementary Figure 8d** and **i**). Regardless, TCF7L2 OE enhanced growth in the WCM1078 model but did not rescue the growth delay induced by A947- and AU-15330-treatment like empty vector (EV) control or β-Catenin OE conditions (**Supplementary Figure 8j**). The fact that forced TCF7L2 expression cannot rescue the phenotype is not surprising, since multiple TCF7L2 binding sites are closing upon SMARCA2/4 degradation making TCF7L2 interaction with these DNA domains impossible.

This let us hypothesize that CRPC-WNT is not driven by the canonical WNT pathway and that TCF7L2 is hijacked to activate other pathways. Although these findings were unexpected, this data aligns with the fact that we do not see any canonical WNT gene sets affected by A947-treatment, despite seeing multiple WNT TFs being downregulated at the protein level over time (**Supplementary Figure 6c** and **8f**). Thus, we raised the question if TCF binding sites have been reprogrammed in CRPC-WNT to drive WNT-independent pathways. Therefore, we aimed to uncover the TCF7L2 orchestrated pathways, which are affected by A947-treatment in CRPC-WNT by performing TCF7L2 ChIP-seq.

### SWI/SNF ATPase degradation abrogates proliferative signaling pathways tied to TCF7L2 in CRPC-WNT

To define the TCF7L2 cistrome in CRPC-WNT, we used chromatin immunoprecipitation followed by sequencing (ChIP-seq) analysis of WCM1078 organoids. In line with the chromatin closure at TCF7L2 motif sites by ATAC-seq, we found decreased TCF7L2 binding to chromatin in WCM1078 organoids upon exposure to A947 for 4h (**Fig. 4a**). A947 treatment led to the loss of 4,393 sites compared with the A858 control. Of the 4,393 lost TCF7L2 sites, 1,903 showed overlap with closing chromatin regions detected by ATAC-seq, representing a significant proportion of downregulated ChIP-seq (43%) and downregulated ATAC-seq peaks (48%). As expected, the top depleted motifs upon A947-treatment in the TCF7L2 ChIP-seq are associated with LEF and TCF7L2 (**Supplementary Figure 10a**). This indicates that SMARCA2/4 degradation indeed interferes with TCF7L2 chromatin binding.

**Fig. 4.**
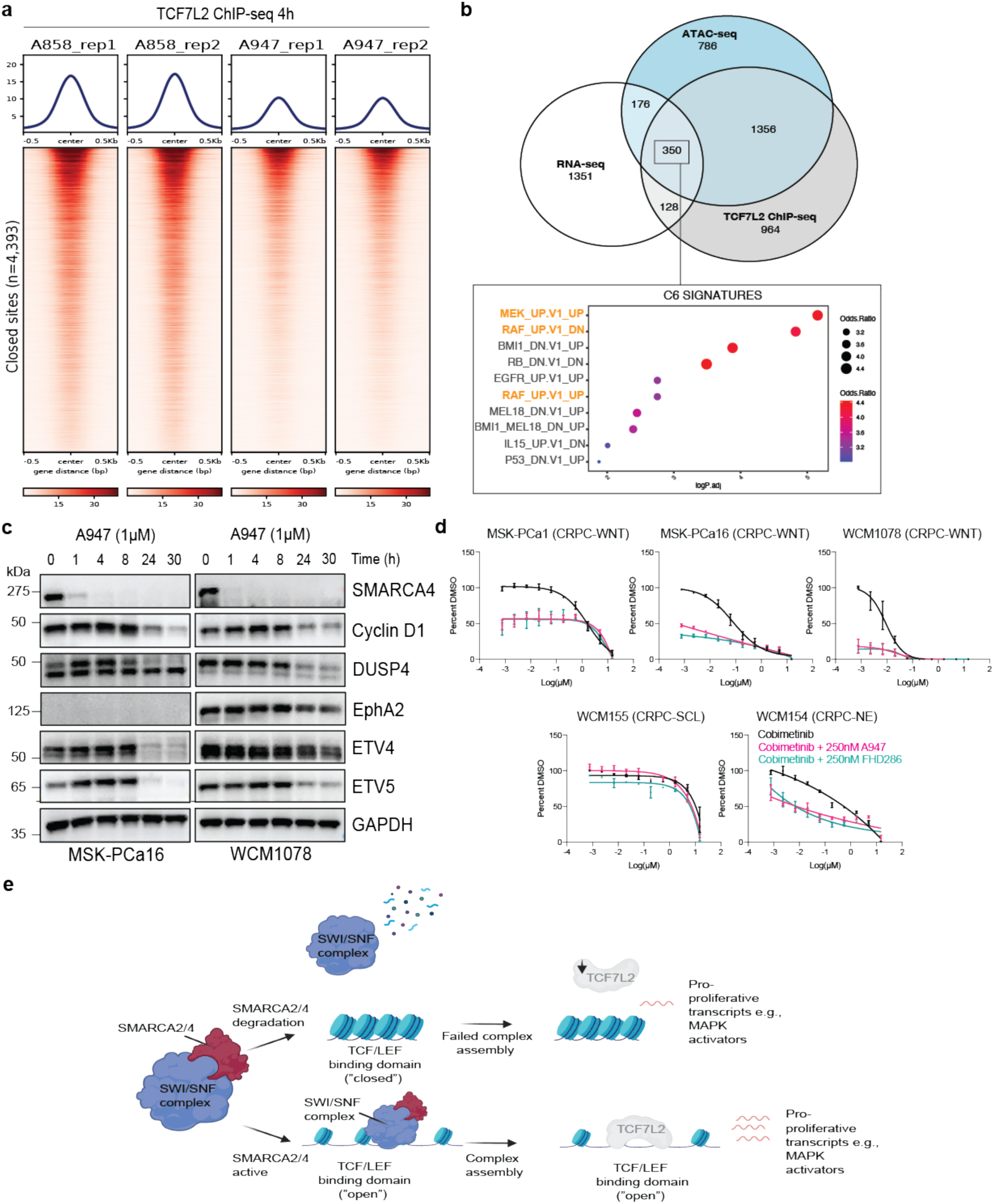
TCF7L2 regulates a pro-proliferative signatures in CRPC-WNT. a, ChIP-seq read-density tornado plots from WCM1078 organoids treated with 1µM A858 or 1µM A947 for 4h (*n* = 2 biological replicates). Venn diagram indicating lost regions. b, Venn diagram indicating A947-treated lost regions from ChIP-seq, ATAC-seq and RNA-seq data in WCM1078. GSEA analysis was performed from 350 overlapping genes. c, Immunoblot of indicated proteins at indicated time upon treatment with 1µM A947. GAPDH serves as loading control. Data is representative of *n* = 2 independent experiments. d, Dose-response curves with indicated drugs after measurement of proliferation with Celltiter Glo 2.0 after 7-day treatment (*n*=2 independent experiments). e, Model of mechanism. SMARCA4-containing SWI/SNF complex maintains open chromatin confirmation of TCF binding domains in CRPC-WNT. TCF7L2 positively regulates lineage-specific programs that orchestrate CRPC-WNT proliferation and survival. TFC7L2 cannot bind to the chromatin when SWISNF ATPase activity is impaired. In addition, loss of SWI/SNF ATPases leads to downregulation of TCF7L2 transcript protein levels. Figure created with Biorender.com.

Gene-set enrichment analysis (GSEA) of the 1903 genes overlapping between ATAC-seq and TCF7L2 ChIP-seq revealed enrichment of depleted peaks in regions associated with proliferative genes, including previously identified MAPK-associated pathways (RAF_UP.V1_DN, EGFR_UP.V1_UP, RAF_UP.V1_UP, MEK_UP.V1_UP) (**Supplementary Figure 10b**). GSEA of the intersect of RNA-seq, ATAC-seq, and ChIP-seq (350 genes) resulted in the top downregulated pathway being MEK signaling (**Fig. 4b**). To understand if CRPC-WNT is dependent on the MAPK-MEK signaling axis, we checked how A947-treatment transcriptionally affects genes that define the so-called MAPK pathway activity score (MPAS)^51^. MPAS contains a set of MAPK downstream targets that selectively predict sensitivity to MEK inhibitors (MEKi) in multiple cancer types. Interestingly, DNPC has previously been described as sensitive to MEKi and FGFR1 inhibition^1^. Moreover, inhibition of MEK or FGFR1 led to downregulation of the gene transcripts making up the MPAS signature (e.g., ETV4, ETV5, DUSP4, SPRY2) in models of DNPC^1^. Indeed, when checking the expression of MPAS genes from RNA-seq data upon A947-treatment, we found that almost all these transcripts were downregulated in the CRPC-WNT model WCM1078 (**Supplementary Figure 11a**). This was confirmed on the protein level in WCM1078 and MSK-PCa16 CRPC-WNT lines when treating the cells with A947 or AU-15330 (**Fig. 4c, Supplementary Figure 11b**). Since the downregulation of MPAS proteins happened only after 24h of A947 treatment, we postulate this effect is downstream of TCF7L2 downregulation (which already happens within 1h) (**Fig. 3e**). ChIP- seq of TCF7L2 confirmed binding of promoter regions of MPAS genes (**Supplementary Figure 12a-d)**. Further, we found that TCF7L2 also binds the promotor of SMARCA4 but not SMARCA2 or PBRM1. This binding is reduced upon the treatment with A947 (**Supplementary Figure 12e-g**). We also found reduced binding at the SIX2 promotor in the A947-treatment condition (**Supplementary Figure 12h**). SIX2 is known to be a TCF7L2 interactor and has implications in PCa lineage identity ^23,52^. Lastly, we tested if the MPAS is indeed predictive for sensitivity to MEK inhibition in CRPC-WNT. For this, we used MEK inhibitor Cobimetinib alone or in combination with A947 or SMARCA2/4 inhibitor FHD-286 in CRPC-WNT, CRPC-NE, and CRPC-SCL lines. We found that Cobimetinib alone and in combination with SMARCA2/4 interfering agents was most active in CRPC-WNT **(Fig. 4d).** These results were recapitulated with another MEK inhibitor, Trametinib **(Supplementary Figure 11c).**

Thus, we conclude that the SWI/SNF complex directly shapes the cistrome for WNT signaling transcription factor TCF7L2 in AR-negative CRPC-WNT to drive pro-proliferative pathways that are predictive of MEK inhibitor sensitivity.

## Discussion

The standard approach to treating advanced prostate cancer has been to modulate the AR axis either through direct or indirect means^53^. Drugs such as enzalutamide or abiraterone are potent androgen- receptor signaling inhibitors (ARSi) used clinically, and other agents, including AR-degraders, are in clinical development. Resistance to ARSi therapy manifests in manifold ways (e.g., *AR* gene mutation, amplification, enhancer amplification), as well as a subset acquiring epigenetic rewiring towards AR- negative phenotypes^7^.

As mentioned, AR-negative PCa had previously been classified as CRPC-NE or DNPC. CRPC-NE is characterized by a small-cell morphology, stemness, and the expression of neuronal and NE marker genes; however, while DPNC is also AR-negative, it shows no evidence of neuroendocrine differentiation based on morphology or expression of classical NE markers^1,54^. From this classification emerged two novel subtypes that branch into the DNPC category: CRPC-WNT and CRPC-SCL^23^. Tang et al. suggested that CRPC-WNT is TCF/LEF TF driven, while CRPC-SCL is YAP/TAZ TF dependent. Unfortunately, targeting these specific pathways directly remains a clinical challenge^55,56^. An alternative approach is to interrupt master transcriptional lineage programs by targeting TF cofactors and associated epigenetic regulators. Among these epigenetic regulators is the chromatin remodeler SWI/SNF complex, which we have previously found to be dysregulated in PCa throughout disease progression and thus represents a viable therapeutic target in early but also late-stage disease^6^. The SWI/SNF complex has been linked to being a predominant orchestrator of lineage-defining transcriptional programs, especially in master TF-addicted cancers^57^.

A recent study in AR-dependent PCa found that PROTAC degraders that target the SWI/SNF complex disrupt the enhancer and promoter looping interaction that wire supra-physiological expression of lineage-driving oncogenes, including the AR, FOXA1, and MYC^11^. However, the number of AR-negative models tested in this study were limited; therefore, we examined the effect of SWI/SNF ATPase PROTAC degraders in a PCa-focused screen. We utilized both AR-dependent and a broad spectrum of AR- negative PCa model systems, including CRPC-NE, CRPC-WNT, and CRPC-SCL. Here, we report that VHL-dependent degraders for SWI/SNF ATPase components decrease proliferation and spheroid formation in organoids of the CRPC-WNT phenotype for which no standard-of-care treatment exists. Clinically, CRPC-WNT tumors account for around 10-11% of all CRPC cases (**Fig. 1i**)^9,11,23^. We found that the SWI/SNF ATPase SMARCA4, but not SMARCA2, is a dependency in the CRPC-WNT phenotype in vitro and in vivo.

Mechanistically, we identified that the activity of intestinal stem cell factor TCF7L2, the most active TF in CRPC-WNT^23^, to be attenuated upon degradation of SMARCA2/4. To our surprise CRPC-WNT models did not respond to classical ways of WNT inhibition. This indicates that TCF7L2 is involved in maintaining a niche of DNPC but potentially via non-canonical, “nontraditional” roles of TCF/LEF signaling. In line with this, we discovered that A947-treatment reduces TCF7L2 binding to MAPK-associated gene promotors. This indicates that TCF7L2 potentially gets hijacked from its traditional role in canonical WNT signaling to assist in driving MAPK transcriptional circuits. In line with these findings canonical WNT signaling has not been nominated as driver of DNPC, emphasizing that TCF7L2 has different roles in the DNPC subtype termed CRPC-WNT^1^. Further, a link between reported Ras pathway activation and TCF7L2 has been reported^58^. This is underpinned by the finding that MAPK signaling is a dependency in DNPC and that clinical trials with MEKi Trametinib in CRPC have entered Phase II (**NCT02881242**)^1,59^. However, these trials were not biomarker-based and were conducted in patients who progressed after AR-targeted therapy. Thus, based on our data, it may be beneficial to clinically assess the utility of the CRPC-WNT score as a biomarker in CRPC to predict response to SMARCA4 or MAPK targeting therapies.

In summary, we nominated the SWI/SNF chromatin remodeling complex, primarily SMARCA4, as a vulnerability in DNPC classified as CRPC-WNT. Impaired maintenance of chromatin accessibility by SMARCA4-containing SWI/SNF complexes potentially blocks the binding of TCF7L2 on the chromatin leading to reduced pro-proliferative pathway activity (**Fig. 4e**). Paralleling other studies in CRPC and small-cell lung cancer (SCLC), our data suggests that SWI/SNF-targeting agents have general efficacy in cancers that are strongly driven by nuanced master transcriptional regulators^11,60^. Further, we posit that MEK inhibition could be another viable approach to target CRPC-WNT and potentially other DNPC subtypes and anticipate a mechanistic connection in future work, as indicated in previous studies^1,2^. We recognize, more in-depth mechanistic studies need to be conducted in this PCa phenotype to fully understand the underlying complex role of TCF7L2.

## Methods

### Cell lines and compounds

PCa cell lines (LNCaP, 22Rv1, VCaP, PC3, DU145, NCI-H660, C4-2), other cell lines (HEK293T, DLD1) and benign prostate line (RWPE-1) were purchased from ATCC and maintained according to ATCC protocols. Patient-derived CRPC organoids (WCM and MSK) were established and maintained as organoids in Matrigel drops according to the previously described protocol^61^. LNCaP-AR cells were a kind gift from Dr. Sawyers and Dr. Mu (Memorial Sloan Kettering Cancer Center) and were cultured as previously described^62^. All used cell lines and their phenotype are listed in Supplementary Table 1. Cell cultures were regularly tested for *Mycoplasma* contamination and confirmed to be negative. Genentech Inc. synthesized A947, its epimer (A858), FHD-286 and AU-15330. Cobimetinib, Trametinib, VL285, MBAS and CHIR99021 were purchased from SelleckChem. BRM014, LGK974 and iCRT14 were purchased from MedChemExpress. All drugs used in this study are listed in Supplementary Table 2.

### Western blot

Whole-cell lysates were prepared in 1x Cell lysis buffer (CST, 9803) supplemented with protease and phosphatase inhibitor cocktail (Thermo Fisher, 78440), and total protein was measured by Pierce BCA Protein Assay Kit (ThermoFisher Scientific, 23225). An equal amount of protein was loaded in SureBlot 10% or 4 to 15%, Bis-Tris Protein Gel (GenScript), and blotted. Subsequently, the nitrocellulose membrane was incubated with primary antibodies overnight in a cold room, shaking. Following incubation with HRP-conjugated secondary antibodies, membranes were imaged on a Vilber Fusion FX imager. Antibodies are listed in Supplementary Table 3.

### Xenograft experiment and pathological assessment

#### Mice

Male NSG (NOD.Cg-Prkdcscid Il2rgtm1Wjl/Sz) mice at the age of 3-5 weeks were purchased from Charles River laboratories.

Mice were allowed to acclimate for 2 weeks before being used for experiments.

All animal studies were approved by the Cantonal Veterinary Ethical Committee, Switzerland (license BE35/2024). Animals were housed in ventilated cages with unrestricted access to presterilized food and fresh water. A maximum of five animals were maintained per cage on Aspen bedding. The ambient temperature was 20^°^C ± 2^°^C, kept at a constant humidity of 50% ± 10%, and on a 12- hour automatic light–dark cycle.

#### A947 administration

Animal was restrained in injection cone for procedure and tail was warmed in 37°C autoclaved water following disinfection of the tail before injection. Vein was visualized by slightly rotation of the tail. Injection was done with 30-gauge needle into one of the lateral veins of the mouse tail. Injection (5ul/g) was processed slowly without aspiration.

After withdrawal of the needle, injection site was carefully compressed with sterile tissue to stop eventual bleeding. Animal was checked and returned to its home cage afterwards.

A947 (40mg/kg) compound was prepared sterile in 10% Hydroxypropyl-β-cyclodextrin and 50mM sodium acetate in water (pH 4.0) freshly on the day of injection.

#### In vivo experimental design

Tumor fragments (from organoid line WCM1078) were transplanted into NSG mice.

Tumor reached measurable size (60-80mm^3^) after 16 days and mice were treated one-time with A947 compound or vehicle intravenously into the lateral tail vein on day 19.

Mice were monitored 3x times per week as well as tumor size was evaluated by digital calipering.

The volume of the tumors was calculated using the formula 4/3pi*((sqrt(L*W))/2)3, where L is the minor tumor axis and W is the major tumor axis. The maximal subcutaneous tumor size/burden allowed (1000 mm^3^) was not exceeded in this study. Tumors and organs were harvested freshly 21 days after treatment. Fresh tissue was snap-frozen and an additional tissue sample was fixed in Formalin (10%) for paraffin embedding. Paraffin embedded tissue was cut and stained with Hematoxylin and eosin stain for blinded histopathologic assessment by a board-certified veterinary pathologist (S.d.B). Throughout the study one animal had to be excluded due to a bacterial infection.

### Immunohistochemistry

Matrigel-extracted organoids were air-dried and subsequently baked at 62 °C for 25min. Immunohistochemistry (IHC) was performed on sections of formalin-fixed paraffin-embedded organoids (FFPE) using a Bond automated immunostainer and the Bond Polymer Refine detection system (Leica Microsystems, IL, USA) by the Translational Research Unit (TRU) platform, Bern. The TCF7L2 antibody (Cell Signaling Technologies, cat# 2569) was used for staining. The intensity of nuclear immunostaining was evaluated on whole slide tissue sections by a pathologist (S.d.B.) blinded to additional pathological and clinical data.

### Cell viability assay

Cells and organoids were plated in 2D onto coated 96-well plates in their respective culture medium and incubated at 37 °C in an atmosphere of 5% CO2. After overnight incubation, a serial dilution of compounds was prepared and added to the plate. The cells were further incubated for 7 days, and the CellTiter-Glo assay (Promega) was then performed according to the manufacturer’s instructions to determine cell proliferation. The luminescence signal from each well was acquired using the Varioskan LUX Plate Reader (Thermo Fisher), and the data were analyzed using GraphPad Prism software (GraphPad Software).

### Classification of CRPC subtypes from publicly available tumor data

Raw data in FASTQ format were obtained from the respective sources (Table 1.) and aligned against the latest human (GRCh38) genome assembly release. Per-sample alignment and generation of feature-barcode matrices were carried out using the STAR-solo algorithm (STAR version 2.7.10b), tailored to the specific sequencing chemistry and the length of cell^63^ barcodes and unique molecular identifiers (UMIs) on a case-by-case basis. We then imported the output feature-barcode matrices in an R environment (R version 4.0.2) and created individual Seurat objects for each sample (Seurat package version 4.0.3)^64–66^. A first round of quality filtering was performed through scuttle package (version 1.8.1)^67^ by discarding outlier and low-quality cells after inspection of commonly used cell-level metrics (i.e., library size, UMI counts per cell, features detected per cell, mitochondrial and ribosomal counts ratio). Further doublet estimation and removal through the DoubletFinder prediction tool (version 2.0.3)^68^ allowed us to drop unnecessary confounding technical artifacts. Thus, we merged our polished samples into a unique Seurat object. Seurat global-scaling normalization and log-transformation method were applied to the complete expression matrix, followed by the selection of highly variable features and linear transformation to prepare the data for dimensional reduction. The latter scaling step also allowed us to regress out unwanted sources of heterogeneity, such as mitochondrial contamination and uneven library sizes. Thus, we determined the optimal number of principal components (PCs) as the lower dimension exhibiting a cumulative percentage of variation greater than 90% to be 82. Uniform manifold approximation and projection (UMAP) based on the previously selected PCs was used to reduce dimensionality and visualize the organization and clustering of cells. Specifically, we applied a graph-based unsupervised approach coupled with the Louvain clustering algorithm implemented in Seurat to generate cell clusters. We evaluated several levels of granularity through clustree (package version 0.5.0)^69^ resolution stability analysis to accurately depict the intrinsic cellular heterogeneity. A clustering resolution of 1 was chosen, and markers identification was conducted by taking advantage of a hurdle model designed explicitly for scRNA-seq data and implemented in the MAST statistical framework^70^, followed by a Bonferroni p-value adjustment to correct for multiple testing. Only genes expressed by at least 70% of cells in the cluster, displaying a significant adjusted p-value (pval_adj < 0.05) and a solid logarithmic fold change (log2FC > 1) were deemed as appropriate markers. After marker-based annotation of major cellular populations, we separated our collection, keeping only epithelial and malignant clusters, encompassing healthy/normal specimens of primary and castration-resistant tumors. The subsetted cells were again subjected to normalization and rescaling, and the relative UMAP was generated using 82 PCs, as before. Graph-based clustering was performed, and a granularity resolution of 0.3 was chosen, resulting in 34 clusters. In-depth annotation was assigned through marker identification. Therefore, we focused on determining which clusters could be associated with the activation of specific biological pathways by assessing the enrichment score (through *AddModuleScore* function) of some recently identified signatures^23^ in the CRPC setting, such as AR, neuroendocrine (NE), stem-cell-like (SCL) and WNT- signaling. The Seurat object was then converted into an *anndata* object, and the Scanpy (version 1.9.5)^71^ toolkit was used for visualization.

**Table 1.** Publicly available datasets were analyzed in this study.

| Dataset ID | Number of samples | Sample type | Sequencing platform and chemistry | Reference (DOI) |
| --- | --- | --- | --- | --- |
| GSE137829 | 6 | CRPC | 10x Genomics 3' v2 | 10.1038/s42003-020-01476-1<sup>25</sup> |
| GSE143791 | 9 | CRPC | 10x Genomics 3' v2 | 10.1016/j.ccell.2021.09.005<sup>26</sup> |
| GSE157703 | 2 | Primary tumor | 10x Genomics 3' v3 | 10.1186/s12943-020-01264-9<sup>29</sup> |
| GSE181294 | 35 | Primary tumor and normal adjacent tissue | 10x Genomics 3' v2 | 10.1038/s41467-023-36325-2<sup>28</sup> |
| GSE193337 | 8 | Primary tumor and normal adjacent tissue | 10x Genomics 3' v3 | 10.1186/s12943-022-01597-7<sup>27</sup> |
| GSE210358 | 14 | CRPC | 10x Genomics 3' v3 | 10.1126/science.abn0478<sup>2</sup> |

### Transfection and siRNA-mediated Knockdown

ON-TARGET plus siRNA SMARTpool siRNAs against *SMARCA4 (L-010431-00-0005), SMARCA2 (L- 017253-00-0005), CTNNB1 (L-003482-00-0005), TCF7L2 (L-003816-00-0005) and control (D-001810-10-05)* were purchased from Dharmacon. Reverse Transfection was performed in 6-well plates using the Lipofectamine 3000 reagent (Thermo Fisher Scientific) to the proportions of 2 μL of 20 μM siRNA per well. After overnight incubation, 5,000 cells were seeded triplicates in a clear 96-well plate, and confluence was monitored using IncuCyte S3 for up to 7 days. The remaining cells were harvested for protein extraction 96 h after transfection.

### siRNA rescue experiment

Rescue sequences (Supplementary Table 4) for the siTools against TCF7L2 (pool of 30 siRNAs) were designed and purchased from siTools Biotech. The rescue plasmid for TCF7L2 was synthesized and purchased from Atum Bio. Reverse Transfection of rescue plasmid was performed in 6-well plates using the Lipofectamine 3000 reagent (Thermo Fisher Scientific) to the proportions of 5 μL of 2 μg plasmid per well. After overnight incubation, cells were transfected with siTools TCF7L2 siRNA pool using the Lipofectamine RNAiMAX reagent (Thermo Fisher Scientific) to the proportions of 2 μL of 20nM siRNA per well. After overnight incubation, 5,000 cells were seeded triplicates in a clear 96-well plate, and confluence was monitored using IncuCyte S3 for up to 7 days. The remaining cells were harvested for protein extraction 96 h after Transfection. Sequences of siRNA and rescue sequence are shown Supplementary Table 4.

### Lentiviral transduction of sgRNA into organoids

Organoids were transduced with CRISPRmod CRISPRi All-in-one Lentiviral hEF1a sgRNA against SMARCA4 particles (sgRNA1: VSGH12442-253241676, sgRNA2: VSGH12442-253150712, sgRNA3: VSGH12442-253336788) or control (VSGC12547). Briefly, 5µl of concentrated virus was added to 500000 cells in suspension with TransDux MAX Lentivirus Transduction Reagent (SBI, LV860A-1) and spinfected at 600 x g, 1h, 32°C. Subsequently, they were incubated 4h at 37°C before careful seeding into Matrigel drops (50000 cells per drop) and topped up with appropriate media. Two days after transduction, virus-integrated cells were selected with puromycin. Sequences for sgRNAs are listed in Supplementary Table 6.

### Incucyte growth assays

#### 2D monolayer formation

A total of 5,000 cells per well were seeded in clear 96-well plates. After overnight incubation, compounds were added at indicated concentrations. Plates were read in an IncuCyte S3. Every 6h, phase object confluence (percentage area) for cell growth was measured. Growth curves were visualized using GraphPad.

#### 3D organoid formation

Twenty thousand cells per well were seeded in Matrigel drops using a clear 48-well plate. After 48h incubation, compounds were added at indicated concentrations. Plates were read in an IncuCyte SX5. Every 6h, organoid object count (µm2/image) for organoid formation was measured. Growth curves were visualized using GraphPad.

### TOPFlash reporter assay

WCM1078 cells were transduced with FOPFlash reporter (LTV-0011-4N, LipExoGen) or TOPFlash reporter (LTV-0011-4S, LipExoGen). After selection with Blasticidin cells were transduced with internal control Renilla Luciferase (Rluc) Lentivirus (BPS Biosciences, 79565-G). After selection with G418, cell were transduced with lentivirus to overexpress the empty vector (GeneCopeeia, NEG-LV105), CTNNB1 (GeneCopoeia, CLP-I4822-LV105-200) or TCF7L2 (GeneCopoeia, CLP-I6388-LV105- 200-GS). After selection with Puromycin, the cells were seeded in triplicates (5000 cells/well) in a 96-well plate. 24h later cells were treated with either DMSO, 1µM A947 or 1µM AU-15330. 48h later, TOPFlash Firefly signal and Renilla internal control signal was detected using the Dual-Glo Luciferase Assay system (Promega, E2920). Luminescence was read using the Varioskan LUX plate reader (Thermo Fisher) and relative luminescence was calculated dividing Firefly with the Renilla luciferease signal. Graphs were were visualized using GraphPad.

### Single-cell RNA-sequencing by SORT-seq library generation and analysis

SORT-seq was performed using Single Cell Discoveries (SCD) service. Organoids were treated for 72h with a control epimer (A858) or active compound (A947) at 1 µM, and 1x10e6 cells were harvested in PBS. Harvested cells were stained with 100ng/ml DAPI to stain dead cells. Using a cell sorter (conducted by Flow Cytometry Core, DBMR, Bern) and the recommended settings (Single Cell Discoveries B.V.), DAPI-negative cells were sorted as single cells in 376 wells of four 384-well plates containing immersion oil per condition. Resulting in a theoretical cell number of 1504 cells per condition. All post-harvesting steps were performed at 4°C. Plates were snap-frozen on dry ice for 15 minutes and sent out for sequencing at Single Cell Discoveries B.V.

Data were analyzed using the Seurat package v.4.3.0^72^. Cell QC filtering was done using the following thresholds: nCount > 4000, nFeature > 1000, percent.mito < 25, log10GenesPerUMI > 0.85. Differential gene expression analysis between clusters was done with Seurat::FindAllMarkers. Module scores were generated with Seurat::AddModuleScore. Gene set enrichment analysis was done with the package fgsea v.1.24.0^73^ and the human gene sets from the Molecular Signatures Database (https://www.gsea-msigdb.org). Gene regulatory networks analysis was done with pySCENIC v.0.12.1^74^. Overall analysis was done in R v.4.2.2.

### RNA-seq library generation and processing

For bulk RNA-seq, organoids were treated with A858 or A947 (1µM) for 24h and 48h (3 biological replicates per condition). RNA was extracted using the RNeasy Kit (Qiagen); library generation and subsequent sequencing was performed by the clinical genomics lab (CGL) at the University of Bern.

Sequencing reads were aligned against the human genome hg38 with STAR v.2.7.3a^75^. Gene counts were generated with RSEM v.1.3.2^76^, whose index was generated using the GENCODE v33 primary assembly annotation. Differential gene expression analysis was done with DESeq2 v.1.34.0^77^. Gene set enrichment analysis was done with the package fgsea v.1.20.0^73^ and the human gene sets from the Molecular Signatures Database (https://www.gsea-msigdb.org). Analysis was done in R v.4.1.2.

### TCF7L2 ChIP-seq library generation and processing

For the ChIP-Seq assay, chromatin was prepared from 2 biological replicates of WCM1078 treated with A858 or A947 (1µM) for 4h, and ChIP-Seq assays were then performed using an antibody against TCF7L2 (Cell Signalling, cat# 2569). ChIP-seq sequence data was processed using an ENCODE- DC/chip-seq-pipeline2 -based workflow (https://github.com/ENCODE-DCC/chip-seq-pipeline2). Briefly, fastq files were aligned on the hg38 human genome reference using Bowtie2 (v2.2.6) followed by alignment sorting (samtools v1.7) of resulting bam files with filtering out of unmapped reads and keeping reads with mapping quality higher than 30. Duplicates were removed with Picard’s MarkDuplicates (v1.126) function, followed by indexation of resulting bam files with samtools. For each bam file, genome coverage was computed with bedtools (v2.26.0), followed by the generation of bigwig (wigToBigWig v377) files. Peaks were called with macs2 (v2.2.4) for each treatment sample using a pooled input alignment (.bam file) as control. Downstream analyses were performed with DiffBind v3.11.1 with default parameters, except for summits=250 in dba.count(). dba.contrast() and dba.analyzed() were used to compute significant differential peaks with DESeq2.

### ATAC-seq library generation and processing

ATAC-seq was performed from 50’000 cryo-preserved cells per condition (1µM A858 and 1µM A947, n = 3 biological replicates) treated for 4h and analyzed as described in previous study^78^. Briefly, 50,000 cryo-preserved cells per condition were lysed for 5 minutes on ice and tagmented for 30 minutes at 37°C, followed by DNA isolation. DNA was barcoded and amplified before sequencing.

### PRO-cap library generation and processing

For PRO-cap, approximately 30 million cells were processed per sample as previously described^79,80^. Library preparations for two biological replicates were performed separately. Cells were permeabilized, and run-on reactions were performed. After RNA isolation, two adaptor ligations and reverse transcription were performed with custom adaptors. Between adaptor ligations, cap state selection reactions were carried out using a series of enzymatic steps. RNA washes, phenol:chloroform extractions and ethanol precipitations were conducted between reactions. All steps were performed under RNase-free conditions. Libraries were sequenced on Illumina’s NovaSeq lane following PCR amplification and library clean-up. Raw sequencing data was processed as previously described^81^. Briefly, sequencing data was trimmed with fastp version 0.22.0 and then aligned to the human genome (hg38) concatenated with EBV and human rDNA sequences (GenBank U13369.1) using STAR V2.7.10b. Raw alignments were filtered with samtools version 1.18 and deduplicated using umi_tools version 1.1.2. Alignments were converted to bigwig files using bedtools version 2.30.0 and kentUtils bedGraphToBigWig V2.8. Peaks were called using PINTS version 1.1.6^41^. Divergent peaks not overlapping with TSS +/-500 bp (GENCODE V37) were regarded as candidate enhancer RNAs. GIGGLE is a genomics search engine that identifies and ranks the significance of shared genomic loci between query features and thousands of genome interval files (in our case a database of ChIP-seq experiments). A higher GIGGLE score means a stronger overlap between query features and features from the database (in our case a ChIP-seq experiment from the Cistrome database). Downstream analysis was done in R v.4.2.2. Heatmaps were generated with deepTools v.3.5.0.

Peaks were annotated with HOMER v.4.11 (http://homer.ucsd.edu/). Distal peaks were defined as those peaks in known introns and intergenic regions, and over 2 kb upstream or downstream from known transcription start sites. GIGGLE scores were generated at http://dbtoolkit.cistrome.org. Analysis was done in R v.4.2.2. Heatmaps were generated with deepTools v.3.5.0.

### TCF7L2 enhancer analysis

Hi-C and RNA-seq data for 80 mCRPC biopsies had previously been generated by the Feng lab. Using the gene markers established by Tang et al., we classified these samples into four subtypes: Stem Cell-like (SCL), Neuroendocrine (NE), Androgen Receptor-dependent (AR), and Wnt-signaling dependent (WNT), based on the mean log expression of the designated marker genes. Among the 80 samples, only three—DTB-135-PRO, DTB-218-BL, and DTB-130-BL—fell into the WNT category. Of these, only DTB-135-PRO had a Hi-C cis interaction depth exceeding 1×10⁸, making it the only viable sample for studying enhancer-promoter interactions in WNT signaling. Using the high-quality DTB-135-PRO dataset, we then applied ICE normalization, as previously described by G. Zhao et al., to our Hi-C matrices at a 10 kb resolution. We examined a ±500 kb region surrounding the *TCF7L2* locus and assessed Hi-C contact frequencies within A858 PRO-cap regions scoring above 10. Notably, the *TCF7L2* promoter exhibited the highest contact frequency (10.1) with the suspected enhancer between chr10:113090000-113100000 region in the DTB-135-PRO sample.

## Data availability

RNA-Seq, ChIP-Seq, ATAC-Seq, and PRO-cap data were deposited at the Zenodo database (10.5281/zenodo.15267271).

## Acknowledgments

We thank the Translational Research Unit (TRU) and the Clinical Genomics Lab (CGL) of the University of Bern for their services. Further, we thank Charles Sawyers (Memorial Sloan Kettering) for the LNCaP-AR cell line. We thank Joanna Cyrta (Institut Curie, Paris) for her commentary and suggestions. We are grateful for the project funding by the Bern Centre for Precision Medicine (BCPM) and the Peter and Traudl Engelhorn Foundation. Scientific computing was partly performed on sciCORE - Scientific Computing Center at the University of Basel. Further, we acknowledge Mariana Ricca for her help editing and preparing this manuscript. We thank Michael Berlin at Arvinas Inc. for assistance with chemical synthesis. D.A.Q. acknowledges funding from the Benioff Initiative for Prostate Cancer Research, the Prostate Cancer Foundation, NCI SPORE 1P50CA275741, and Department of Defense awards W81XWH-22-1-0833 and HT94252410252. This work was supported by the Office of the Assistant Secretary of Defense for Health Affairs through the Prostate Cancer Research Program under Award No. HT94252410123.

## Author Contributions

P.T. and M.A.R. conceived the project and designed all studies in the project. P.T., with assistance of I.P., P.D.R., A.N., and L.M., performed most wet-lab experiments. M.S. performed ATAC-seq, B.D.

performed ChIP-seq. A.B., X.Y., and Ju.T. are professional bioinformaticians and conducted data acquisition, analysis, and interpretation of RNA-seq, scRNA-seq, ATAC-seq, ChIP-seq, and PRO-cap.

S.R.S. performed PRO-cap library preparation. A.K.L assisted with PRO-cap bioinformatic analysis. N.L., D.A.Q. and I.P. performed Hi-C enhancer-promoter analysis. M.L. performed animal experiments. S.d.B conducted histopathological evaluation. Jo.T. performed histopathological data analysis. G.C. and M.B. performed bioinformatic CRPC signature score evaluation in scRNA-seq data curated from commercially available data. H.B. and Y.C. provided PCa organoid models. S.P. and C.N. provided expert commentary and bioinformatic expertise. H.Y. provided expert commentary and supervision of PRO-Cap experiment. R.L.Y. provided resources, relevant data, and expert commentary. P.T. and M.A.R. wrote the manuscript with the help of Jo.T.

## References

1. Bluemn, E.G. et al. Androgen Receptor Pathway-Independent Prostate Cancer Is Sustained through FGF Signaling. Cancer Cell 32, 474–489 e6 (2017).

2. Chan, J.M. et al. Lineage plasticity in prostate cancer depends on JAK/STAT inflammatory signaling. Science 377, 1180–1191 (2022).

3. Beltran, H. et al. Divergent clonal evolution of castration-resistant neuroendocrine prostate cancer. Nat Med 22, 298–305 (2016).

4. Ye, Y., Chen, X. & Zhang, W. Mammalian SWI/SNF Chromatin Remodeling Complexes in Embryonic Stem Cells: Regulating the Balance Between Pluripotency and Differentiation. Front Cell Dev Biol 8, 626383 (2020).

5. Hodges, C., Kirkland, J.G. & Crabtree, G.R. The Many Roles of BAF (mSWI/SNF) and PBAF Complexes in Cancer. Cold Spring Harb Perspect Med 6(2016).

6. Cyrta, J. et al. Role of specialized composition of SWI/SNF complexes in prostate cancer lineage plasticity. Nature Communications 11, 5549 (2020).

7. Rubin, M.A., Bristow, R.G., Thienger, P.D., Dive, C. & Imielinski, M. Impact of Lineage Plasticity to and from a Neuroendocrine Phenotype on Progression and Response in Prostate and Lung Cancers. Molecular Cell 80, 562–577 (2020).

8. Xue, Y. et al. SMARCA4 loss is synthetic lethal with CDK4/6 inhibition in non-small cell lung cancer. Nat Commun 10, 557 (2019).

9. Cantley, J. et al. Selective PROTAC-mediated degradation of SMARCA2 is efficacious in SMARCA4 mutant cancers. Nat Commun 13, 6814 (2022).

10. Hoffman, G.R. et al. Functional epigenetics approach identifies BRM/SMARCA2 as a critical synthetic lethal target in BRG1-deficient cancers. Proc Natl Acad Sci U S A 111, 3128–33 (2014).

11. Xiao, L. et al. Targeting SWI/SNF ATPases in enhancer-addicted prostate cancer. Nature 601, 434–439 (2022).

12. Griffin, C.T., Curtis, C.D., Davis, R.B., Muthukumar, V. & Magnuson, T. The chromatin-remodeling enzyme BRG1 modulates vascular Wnt signaling at two levels. Proc Natl Acad Sci U S A 108, 2282–7 (2011).

13. Park, J.I. et al. Telomerase modulates Wnt signalling by association with target gene chromatin. Nature 460, 66–72 (2009).

14. Wang, Y. et al. The long noncoding RNA lncTCF7 promotes self-renewal of human liver cancer stem cells through activation of Wnt signaling. Cell Stem Cell 16, 413–25 (2015).

15. Murillo-Garzon, V. & Kypta, R. WNT signalling in prostate cancer. Nat Rev Urol 14, 683–696 (2017).

16. Miyamoto, D.T. et al. RNA-Seq of single prostate CTCs implicates noncanonical Wnt signaling in antiandrogen resistance. Science 349, 1351–6 (2015).

17. Korinek, V. et al. Constitutive transcriptional activation by a beta-catenin-Tcf complex in APC-/- colon carcinoma. Science 275, 1784–7 (1997).

18. Zhan, T., Rindtorff, N. & Boutros, M. Wnt signaling in cancer. Oncogene 36, 1461–1473 (2017).

19. Sandsmark, E. et al. A novel non-canonical Wnt signature for prostate cancer aggressiveness. Oncotarget 8, 9572–9586 (2017).

20. Grasso, C.S. et al. The mutational landscape of lethal castration-resistant prostate cancer. Nature 487, 239–43 (2012).

21. Zong, Y. et al. Stromal epigenetic dysregulation is sufficient to initiate mouse prostate cancer via paracrine Wnt signaling. Proc Natl Acad Sci U S A 109, E3395–404 (2012).

22. Li, X. et al. Prostate tumor progression is mediated by a paracrine TGF-beta/Wnt3a signaling axis. Oncogene 27, 7118–30 (2008).

23. Tang, F. et al. Chromatin profiles classify castration-resistant prostate cancers suggesting therapeutic targets. Science 376, eabe1505 (2022).

24. Ding, Y. et al. Chromatin remodeling ATPase BRG1 and PTEN are synthetic lethal in prostate cancer. The Journal of Clinical Investigation 129, 759–773 (2019).

25. Dong, B. et al. Single-cell analysis supports a luminal-neuroendocrine transdifferentiation in human prostate cancer. Commun Biol 3, 778 (2020).

26. Kfoury, Y. et al. Human prostate cancer bone metastases have an actionable immunosuppressive microenvironment. Cancer Cell 39, 1464–1478 e8 (2021).

27. Heidegger, I. et al. Comprehensive characterization of the prostate tumor microenvironment identifies CXCR4/CXCL12 crosstalk as a novel antiangiogenic therapeutic target in prostate cancer. Mol Cancer 21, 132 (2022).

28. Hirz, T. et al. Dissecting the immune suppressive human prostate tumor microenvironment via integrated single-cell and spatial transcriptomic analyses. Nat Commun 14, 663 (2023).

29. Ma, X. et al. Identification of a distinct luminal subgroup diagnosing and stratifying early stage prostate cancer by tissue-based single-cell RNA sequencing. Mol Cancer 19, 147 (2020).

30. Muraro, M.J. et al. A Single-Cell Transcriptome Atlas of the Human Pancreas. Cell Syst 3, 385–394 e3 (2016).

31. Aibar, S. et al. SCENIC: single-cell regulatory network inference and clustering. Nat Methods 14, 1083–1086 (2017).

32. Gan, X.Q. et al. Nuclear Dvl, c-Jun, beta-catenin, and TCF form a complex leading to stabilization of beta-catenin-TCF interaction. J Cell Biol 180, 1087–100 (2008).

33. Toualbi, K. et al. Physical and functional cooperation between AP-1 and beta-catenin for the regulation of TCF-dependent genes. Oncogene 26, 3492–502 (2007).

34. Nateri, A.S., Spencer-Dene, B. & Behrens, A. Interaction of phosphorylated c-Jun with TCF4 regulates intestinal cancer development. Nature 437, 281–5 (2005).

35. Guo, Q. et al. A beta-catenin-driven switch in TCF/LEF transcription factor binding to DNA target sites promotes commitment of mammalian nephron progenitor cells. Elife 10(2021).

36. Park, J.S. et al. Six2 and Wnt regulate self-renewal and commitment of nephron progenitors through shared gene regulatory networks. Dev Cell 23, 637–51 (2012).

37. Leppanen, N. et al. SIX2 promotes cell plasticity via Wnt/beta-catenin signalling in androgen receptor independent prostate cancer. Nucleic Acids Res 52, 5610–5623 (2024).

38. Tierney, M.T. et al. Vitamin A resolves lineage plasticity to orchestrate stem cell lineage choices. Science 383, eadi7342 (2024).

39. Schick, S. et al. Acute BAF perturbation causes immediate changes in chromatin accessibility. Nat Genet 53, 269–278 (2021).

40. Iurlaro, M. et al. Mammalian SWI/SNF continuously restores local accessibility to chromatin. Nat Genet 53, 279–287 (2021).

41. Yao, L. et al. A comparison of experimental assays and analytical methods for genome-wide identification of active enhancers. Nat Biotechnol 40, 1056–1065 (2022).

42. Tippens, N.D. et al. Transcription imparts architecture, function and logic to enhancer units. Nat Genet 52, 1067–1075 (2020).

43. Layer, R.M. et al. GIGGLE: a search engine for large-scale integrated genome analysis. Nat Methods 15, 123–126 (2018).

44. Zhao, S.G. et al. Integrated analyses highlight interactions between the three-dimensional genome and DNA, RNA and epigenomic alterations in metastatic prostate cancer. Nat Genet 56, 1689–1700 (2024).

45. Savic, D. et al. Alterations in TCF7L2 expression define its role as a key regulator of glucose metabolism. Genome Res 21, 1417–25 (2011).

46. Savic, D., Park, S.Y., Bailey, K.A., Bell, G.I. & Nobrega, M.A. In vitro scan for enhancers at the TCF7L2 locus. Diabetologia 56, 121–5 (2013).

47. Veeman, M.T., Slusarski, D.C., Kaykas, A., Louie, S.H. & Moon, R.T. Zebrafish prickle, a modulator of noncanonical Wnt/Fz signaling, regulates gastrulation movements. Curr Biol 13, 680–5 (2003).

48. Liu, J. et al. Targeting Wnt-driven cancer through the inhibition of Porcupine by LGK974. Proc Natl Acad Sci U S A 110, 20224–9 (2013).

49. Gonsalves, F.C. et al. An RNAi-based chemical genetic screen identifies three small-molecule inhibitors of the Wnt/wingless signaling pathway. Proc Natl Acad Sci U S A 108, 5954–63 (2011).

50. Hwang, S.Y. et al. Direct Targeting of beta-Catenin by a Small Molecule Stimulates Proteasomal Degradation and Suppresses Oncogenic Wnt/beta-Catenin Signaling. Cell Rep 16, 28–36 (2016).

51. Wagle, M.C. et al. A transcriptional MAPK Pathway Activity Score (MPAS) is a clinically relevant biomarker in multiple cancer types. NPJ Precis Oncol 2, 7 (2018).

52. Shtutman, M. et al. The cyclin D1 gene is a target of the beta-catenin/LEF-1 pathway. Proc Natl Acad Sci U S A 96, 5522–7 (1999).

53. Labbe, D.P. & Brown, M. Transcriptional Regulation in Prostate Cancer. Cold Spring Harb Perspect Med 8(2018).

54. Yamada, Y. & Beltran, H. Clinical and Biological Features of Neuroendocrine Prostate Cancer. Curr Oncol Rep 23, 15 (2021).

55. Neiheisel, A., Kaur, M., Ma, N., Havard, P. & Shenoy, A.K. Wnt pathway modulators in cancer therapeutics: An update on completed and ongoing clinical trials. Int J Cancer 150, 727–740 (2022).

56. Luo, M. et al. Advances of targeting the YAP/TAZ-TEAD complex in the hippo pathway for the treatment of cancers. Eur J Med Chem 244, 114847 (2022).

57. Centore, R.C. et al. Pharmacologic inhibition of BAF chromatin remodeling complexes as a therapeutic approach to transcription factor-dependent cancers. bioRxiv, 2023.09.11.557162 (2023).

58. Kundu, S. et al. Linking FOXO3, NCOA3, and TCF7L2 to Ras pathway phenotypes through a genome-wide forward genetic screen in human colorectal cancer cells. Genome Med 10, 2 (2018).

59. Nickols, N.G. et al. MEK-ERK signaling is a therapeutic target in metastatic castration resistant prostate cancer. Prostate Cancer Prostatic Dis 22, 531–538 (2019).

60. He, T. et al. Targeting the mSWI/SNF Complex in POU2F-POU2AF Transcription Factor-Driven Malignancies. bioRxiv (2024).

61. Gao, D. et al. Organoid cultures derived from patients with advanced prostate cancer. Cell 159, 176–187 (2014).

62. Mu, P. et al. SOX2 promotes lineage plasticity and antiandrogen resistance in TP53- and RB1- deficient prostate cancer. Science 355, 84–88 (2017).

63. Kaminow, B., Yunusov, D. & Dobin, A. STARsolo: accurate, fast and versatile mapping/quantification of single-cell and single-nucleus RNA-seq data. bioRxiv, 2021.05.05.442755 (2021).

64. Stuart, T. et al. Comprehensive Integration of Single-Cell Data. Cell 177, 1888–1902 e21 (2019).

65. Butler, A., Hoffman, P., Smibert, P., Papalexi, E. & Satija, R. Integrating single-cell transcriptomic data across different conditions, technologies, and species. Nat Biotechnol 36, 411–420 (2018).

66. Satija, R., Farrell, J.A., Gennert, D., Schier, A.F. & Regev, A. Spatial reconstruction of single-cell gene expression data. Nat Biotechnol 33, 495–502 (2015).

67. McCarthy, D.J., Campbell, K.R., Lun, A.T. & Wills, Q.F. Scater: pre-processing, quality control, normalization and visualization of single-cell RNA-seq data in R. Bioinformatics 33, 1179–1186 (2017).

68. McGinnis, C.S., Murrow, L.M. & Gartner, Z.J. DoubletFinder: Doublet Detection in Single-Cell RNA Sequencing Data Using Artificial Nearest Neighbors. Cell Syst 8, 329–337 e4 (2019).

69. Zappia, L. & Oshlack, A. Clustering trees: a visualization for evaluating clusterings at multiple resolutions. Gigascience 7(2018).

70. Finak, G. et al. MAST: a flexible statistical framework for assessing transcriptional changes and characterizing heterogeneity in single-cell RNA sequencing data. Genome Biol 16, 278 (2015).

71. Wolf, F.A., Angerer, P. & Theis, F.J. SCANPY: large-scale single-cell gene expression data analysis. Genome Biol 19, 15 (2018).

72. Hao, Y. et al. Integrated analysis of multimodal single-cell data. Cell 184, 3573–3587 e29 (2021).

73. Korotkevich, G. et al. Fast gene set enrichment analysis. bioRxiv, 060012 (2021).

74. Van de Sande, B. et al. A scalable SCENIC workflow for single-cell gene regulatory network analysis. Nat Protoc 15, 2247–2276 (2020).

75. Dobin, A. et al. STAR: ultrafast universal RNA-seq aligner. Bioinformatics 29, 15–21 (2012).

76. Li, B. & Dewey, C.N. RSEM: accurate transcript quantification from RNA-Seq data with or without a reference genome. BMC Bioinformatics 12, 323 (2011).

77. Love, M.I., Huber, W. & Anders, S. Moderated estimation of fold change and dispersion for RNA- seq data with DESeq2. Genome Biol 15, 550 (2014).

78. Hagenbeek, T.J. et al. An allosteric pan-TEAD inhibitor blocks oncogenic YAP/TAZ signaling and overcomes KRAS G12C inhibitor resistance. Nat Cancer 4, 812–828 (2023).

79. Kwak, H., Fuda, N.J., Core, L.J. & Lis, J.T. Precise maps of RNA polymerase reveal how promoters direct initiation and pausing. Science 339, 950–3 (2013).

80. Mahat, D.B. et al. Base-pair-resolution genome-wide mapping of active RNA polymerases using precision nuclear run-on (PRO-seq). Nature Protocols 11, 1455–1476 (2016).

81. Cotter, K.A. et al. Capped nascent RNA sequencing reveals novel therapy-responsive enhancers in prostate cancer. bioRxiv, 2022.04.08.487666 (2022).

