## Supplemental Figures for "A double-negative prostate cancer subtype is vulnerable to SWI/SNF-targeting degrader molecules"

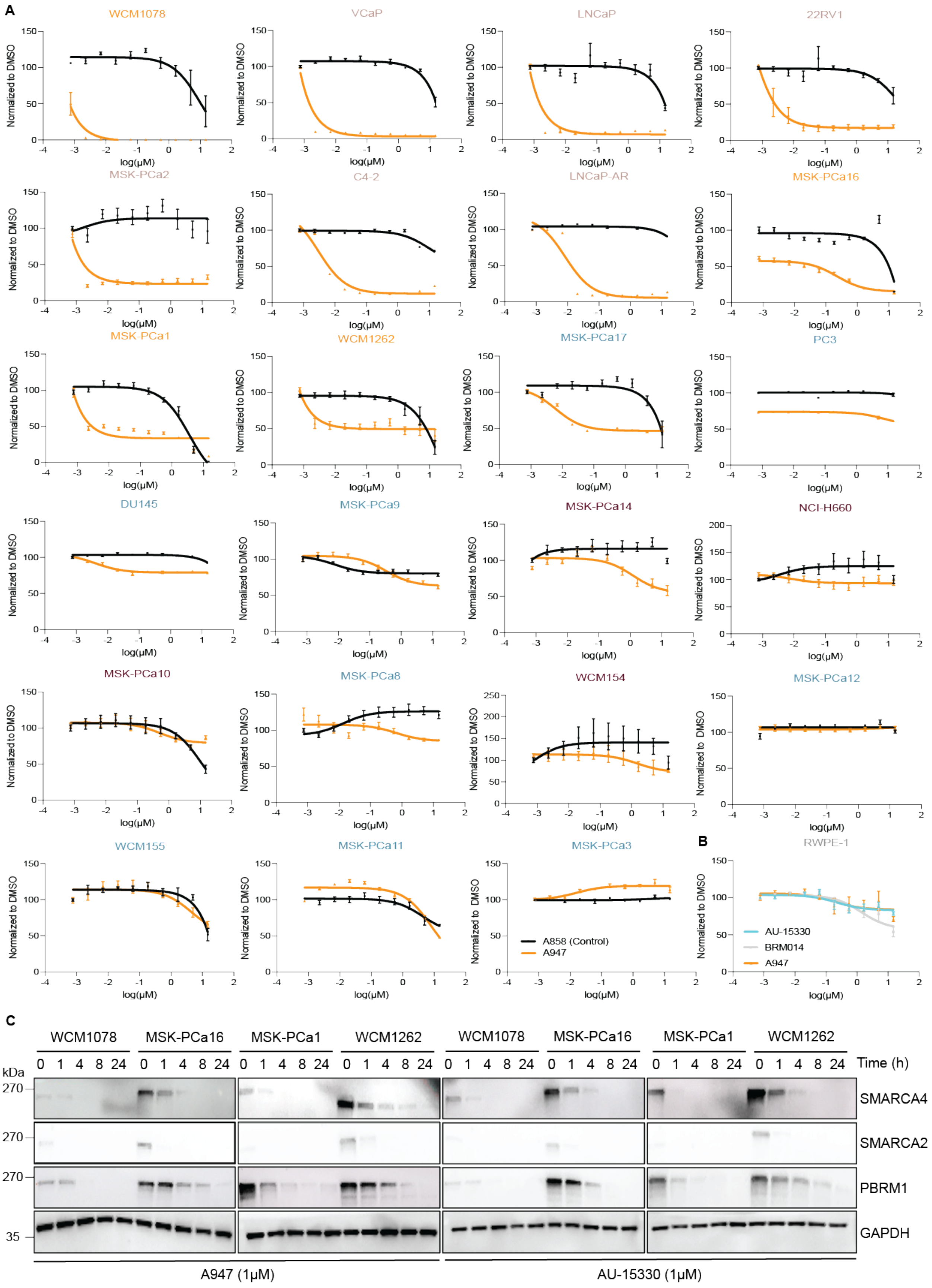


41 **Supplementary Figure 1. Drug screen with SMARCA2/4 degrader**

a, Proliferation measured with CellTiter Glo 2.0 at 7 days treated with dose response of indicated drugs. Data is representative of $n = 3$ independent experiments. Data are presented as mean values +/− SEM.

b, Proliferation measured with CellTiter Glo 3D at 7 days treated with dose response of indicated drugs in benign prostate cell line RWPE-1. Data is representative of $n = 3$ independent experiments. Data are presented as mean values +/− SEM.

c, Immunoblot of indicated proteins in CRPC-WNT cell lines treated with A947 (1μM) or AU-15330 (1μM) for the indicated time. GAPDH serves as a loading control and is probed in a representative immunoblot. Data is representative of $n = 2$ independent experiments.

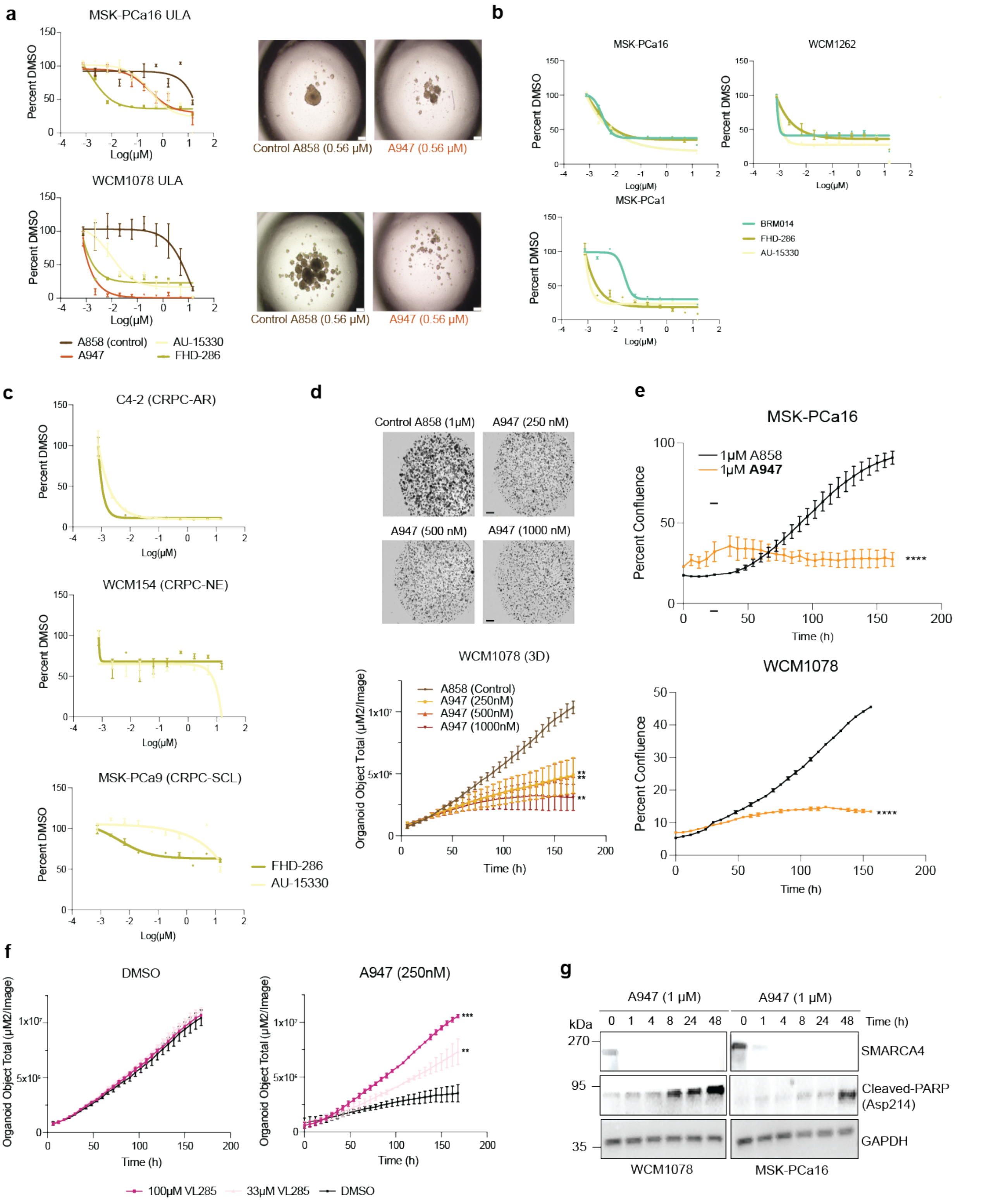


**Supplementary Figure 2. CRPC-WNT is a clinically relevant subtype that can be targeted by SMARCA2/4 PROTAC degraders**

a, Proliferation measured with CellTiter Glo 2.0 at 7 days treated with dose response of indicated drugs in organoids grown in ultra-low attachment (ULA) plates. Data is representative of $n = 2$ independent experiments. Brightfield microscopy pictures were taken 7 days after treatment. Scale bar: 100μm. Data are presented as mean values +/− SEM.

b, Proliferation measured with CellTiter Glo 3D at 7 days treated with dose response of indicated drugs. Data is representative of $n = 3$ independent experiments. Data are presented as mean values +/− SEM.

c, Proliferation measured with CellTiter Glo 3D at 7 days treated with dose response of indicated drugs. Data is representative of $n = 3$ independent experiments. Data are presented as mean values +/− SEM.

d, Spheroid formation measured with Incucyte SX5 upon treatment with indicated doses of A947 and A858. Brightfield microscopy pictures were taken at the endpoint. Scale bar: 800μm. Data is representative of $n = 2$ independent experiments. Data are presented as mean values +/− SEM and analyzed using two-way ANOVA (**$p < 0.01$).

e, Confluence measured with Incucyte S3 upon treatment with indicated doses of A947 and A858. Data is representative of $n = 3$ independent experiments. Data are presented as mean values +/− SEM and analyzed using two-way ANOVA (****$p < 0.0001$).

f, Spheroid formation measured with Incucyte SX5 treated with indicated doses of VL285 or DMSO combined with a dose of 250nM A947. Data is representative of $n = 2$ independent experiments. Error bars are represented as S.E.M. Data are presented as mean values +/− SEM and analyzed using two-way ANOVA ( **$p < 0.01$, ***$p < 0.001$).

g, Immunoblot of indicated proteins in CRPC-WNT cell lines treated with A947 (1μM) for the indicated time. GAPDH serves as a loading control and is probed in a representative immunoblot. Data is representative of $n = 2$ independent experiments.

h, Proliferation measured with CellTiter Glo 3D at 7 days treated with dose response of indicated drugs. Data is representative of $n = 3$ independent experiments. Data are presented as mean values +/− SEM.

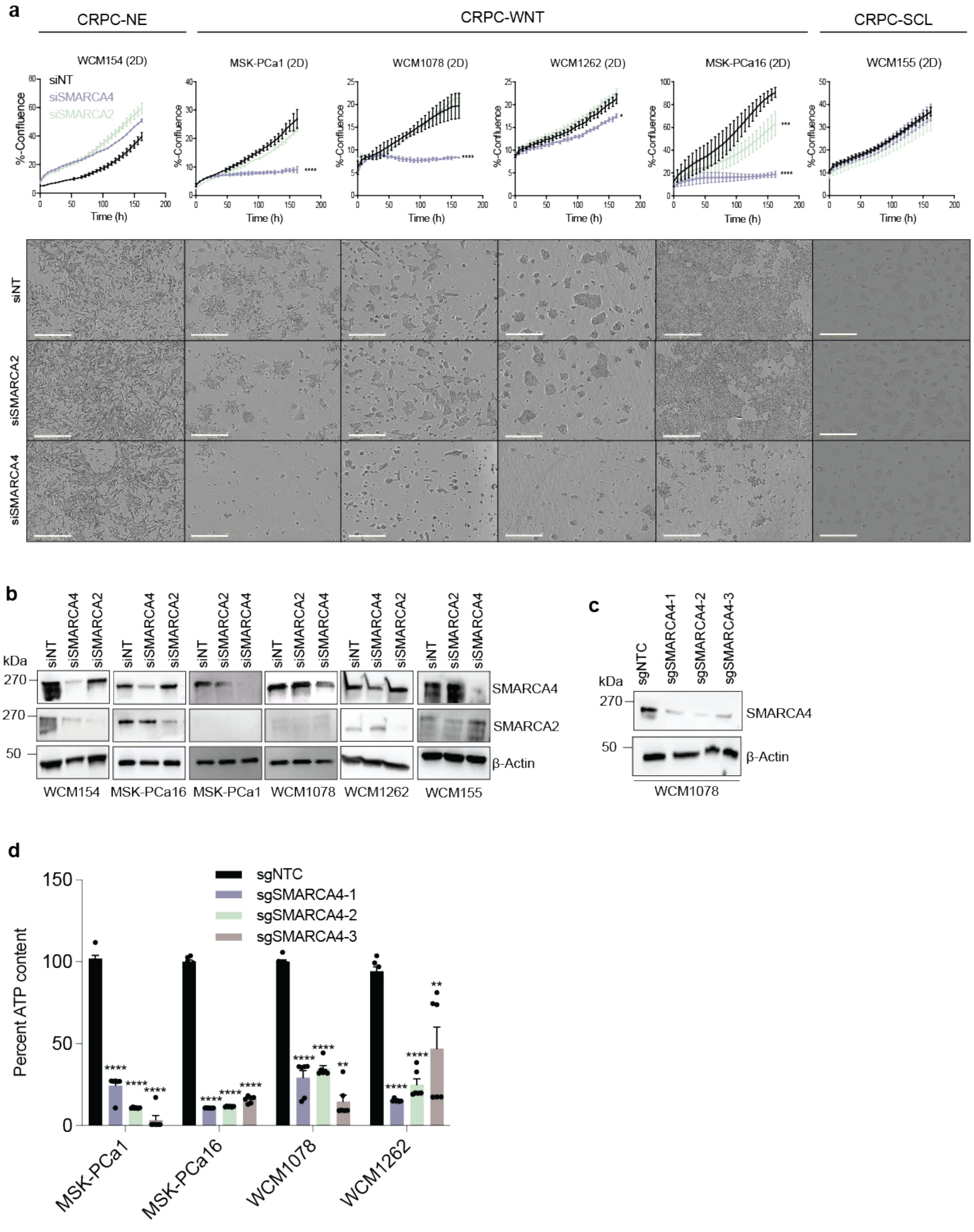
a
CRPC-NE
CRPC-WNT
CRPC-SCL
WCM154 (2D)
MSK-PCa1 (2D)
WCM1078 (2D)
WCM1262 (2D)
MSK-PCa16 (2D)
WCM155 (2D)
siNT
siSMARCA4
siSMARCA2
%-Confluence
Time (h)
siNT
siSMARCA2
siSMARCA4
b
kDa
270
270
50
SMARCA4
SMARCA2
β-Actin
WCM154
MSK-PCa16
MSK-PCa1
WCM1078
WCM1262
WCM155
c
kDa
sgNTC
sgSMARCA4-1
sgSMARCA4-2
sgSMARCA4-3
270
50
SMARCA4
β-Actin
WCM1078
d
sgNTC
sgSMARCA4-1
sgSMARCA4-2
sgSMARCA4-3
Percent ATP content
150
100
50
0
MSK-PCa1
MSK-PCa16
WCM1078
WCM1262

**Supplementary Figure 3. CRPC-WNT organoids are SMARCA4 but not SMARCA2 dependent**

a, Confluence of indicated cell lines transduced with siNon-targeting (NT), siSMARCA2 or siSMARCA4 measured by live-cell imaging using Incucyte S3 ($n = 3$ independent experiments). Brightfield microscopy of indicated cell lines at Incucyte assay endpoint, Scale bars: 400μm. Data are presented as mean values +/− SEM and analyzed using two-way ANOVA (*$p < 0.05$, ***$p < 0.001$, ****$p < 0.0001$).

b, Immunoblot of indicated proteins in indicated cell lines transfected with respective siRNA after 72h. β-Actin represents loading control and is probed in a representative immunoblot. Data is representative of $n = 3$ independent experiments.

c, Immunoblot of indicated proteins in WCM1078 transduced with respective sgRNA after 72h. β-Actin represents loading control and is probed in a representative immunoblot. Data is representative of $n = 2$ independent experiments.

d, Proliferation was measured with CellTiter Glo 3D, and the indicated cell lines were transduced with sgRNAs against SMARCA4 after 10 days of growth ($n = 3$ independent biological experiments).

e, Immunoblot of indicated proteins in indicated cell lines treated with 100nM A947 for up to 48h. GAPDH serves as loading control. Data is representative of $n = 2$ independent experiments. Data are presented as mean values +/− SEM and analyzed using Students t-test (**$p < 0.001$, ***$p < 0.001$, ****$p < 0.0001$).

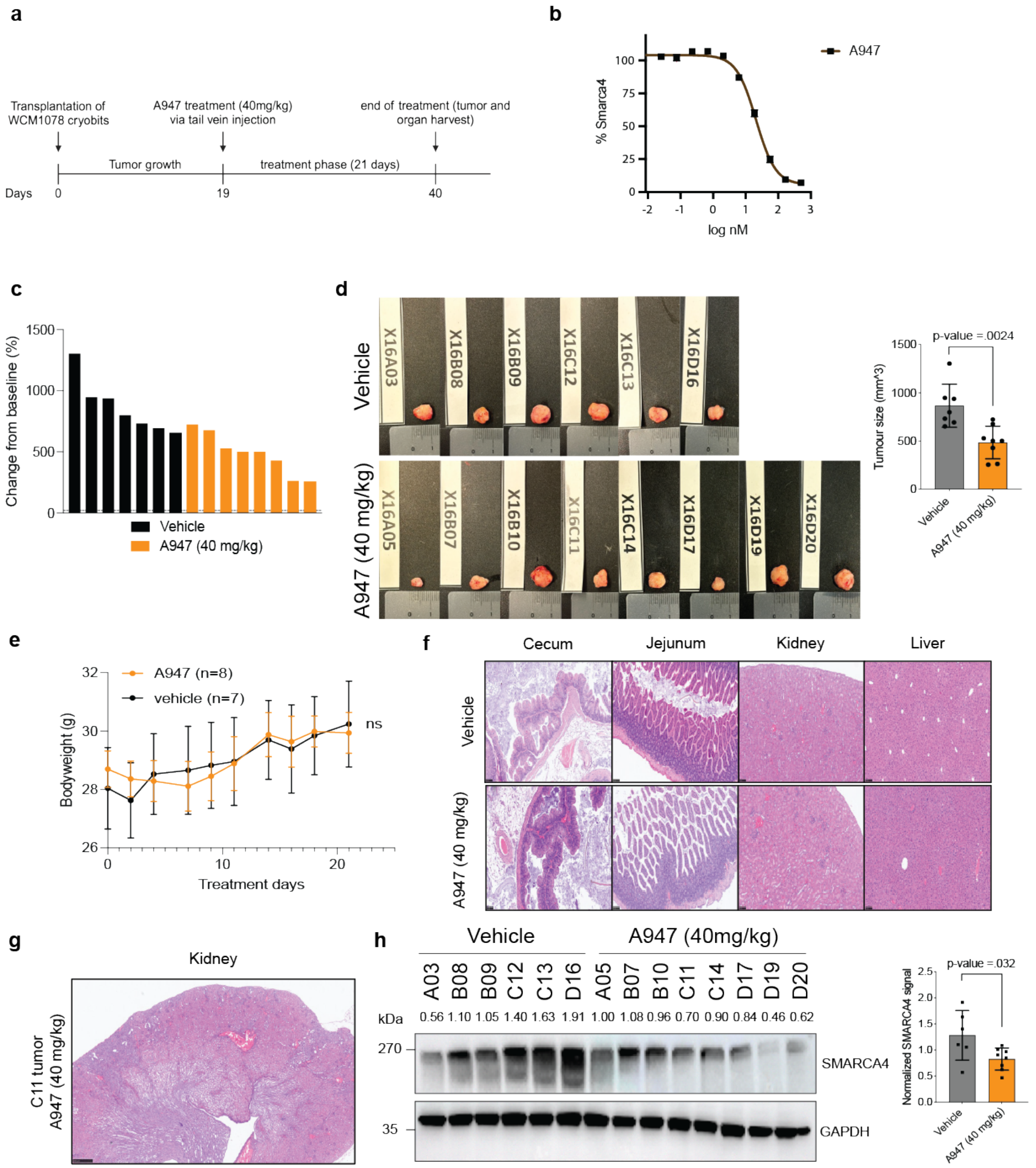
a
Transplantation of WCM1078 cryobits
A947 treatment (40mg/kg) via tail vein injection
end of treatment (tumor and organ harvest)
Tumor growth
treatment phase (21 days)
Days 0 19 40
b
% Smarca4
log nM
A947
c
Change from baseline (%)
Vehicle
A947 (40 mg/kg)
d
Vehicle
A947 (40 mg/kg)
X16A03 X16B08 X16B09 X16C12 X16C13 X16D16
X16A05 X16B07 X16B10 X16C11 X16C14 X16D17 X16D19 X16D20
p-value =.0024
Tumour size (mm^3)
Vehicle
A947 (40 mg/kg)
e
A947 (n=8)
vehicle (n=7)
ns
Bodyweight (g)
Treatment days
f
Cecum
Jejunum
Kidney
Liver
Vehicle
A947 (40 mg/kg)
g
Kidney
C11 tumor
A947 (40 mg/kg)
h
Vehicle
A947 (40mg/kg)
A03 B08 B09 C12 C13 D16 A05 B07 B10 C11 C14 D17 D19 D20
kDa
0.56 1.10 1.05 1.40 1.63 1.91 1.00 1.08 0.96 0.70 0.90 0.84 0.46 0.62
270
35
SMARCA4
GAPDH
p-value =.032
Normalized SMARCA4 signal
Vehicle
A947 (40 mg/kg)

96

97

**Supplementary Figure 4. A947-treatment is reducing CRPC-WNT tumor growth in vivo with minimal adverse effects**

a, Timeline of in vivo experiment. 15 mice were treated for 21 days with a single dose of 40mg/kg A947 once tumors reached a size of at least 60 mm3.

b, Percent of SMARCA4 expression measured by immunofluorescence at indicated doses of A947 in murine lung adenoma LA4 cells. Data are presented as mean values +/− SEM.

c, Waterfall plot showing change from percent baseline of each individual tumor.

d, Pictures of tumors taken after mice were sacrificed and quantification of average tumor size in indicated conditions. Data are presented as mean values +/− SEM and analyzed using Students t-test (p-val=.0024).

e, Average mouse bodyweight (g) over time with indicated treatment. Data are presented as mean values +/− SEM and analyzed using two-way ANOVA (ns=not significant).

f, H&E stainings of indicated tissues taken from mice after endpoint sacrifice. Scale bar: 100µm.

g, H&E staining of kidney in mouse with chromic renal interstitial fibrosis and tubular atrophy taken from mice after endpoint sacrifice. Scale bar: 100µm.

h, Immunoblot of indicated proteins in tumors taken at endpoint of xenograft experiment treated with vehicle or A947. GAPDH represents loading control and is probed in a representative immunoblot. Data is representative of *n* = 2 independent experiments.Numbers above blot are loading control normalized quantification determined using ImageJ, average of normalized signal is represented as a bar chart. Data are presented as mean values +/− SEM and analyzed using Students t-test (p-value=0.032).

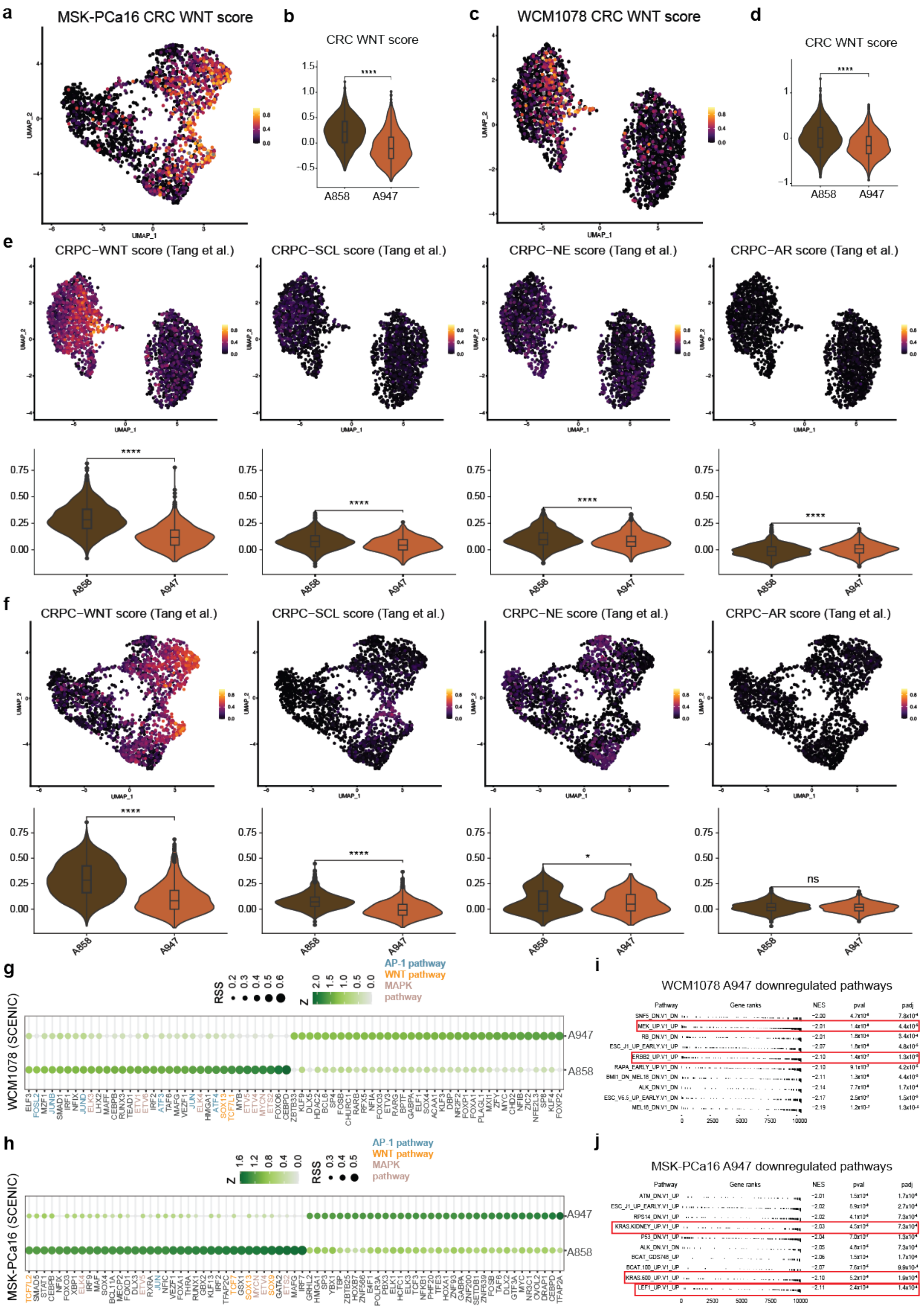
a
MSK-PCa16 CRC WNT score
b
CRC WNT score
c
WCM1078 CRC WNT score
d
CRC WNT score
A858
A947
e
CRPC−WNT score (Tang et al.)
CRPC−SCL score (Tang et al.)
CRPC−NE score (Tang et al.)
CRPC−AR score (Tang et al.)
f
CRPC−WNT score (Tang et al.)
CRPC−SCL score (Tang et al.)
CRPC−NE score (Tang et al.)
CRPC−AR score (Tang et al.)
g
WCM1078 (SCENIC)
AP-1 pathway
WNT pathway
MAPK pathway
RSS
Z
h
MSK-PCa16 (SCENIC)
i
WCM1078 A947 downregulated pathways
j
MSK-PCa16 A947 downregulated pathways
Pathway
Gene ranks
NES
pval
padj

**Supplementary Figure 5. A947-treatment consistently modulates lineage-defining CRPC-WNT signature and master TFs**

a, UMAP plot of MSK-PCa16 organoids treated with either 1μM A858 or 1μM A947 for 72h displaying colorectal WNT (CRC WNT) signature score.

b, Violin plot of MSK-PCa16 organoids treated with either 1μM A858 or 1μM A947 for 72h displaying CRC WNT signature score. Analyzed using the Wilcoxon test (****$p < 0.0001$).

c, UMAP plot of WCM1078organoids treated with either 1μM A858 or 1μM A947 for 72h displaying CRC WNT signature score.

d, Violin plot of WCM1078 organoids treated with either 1μM A858 or 1μM A947 for 72h displaying CRC WNT signature score. Analyzed using the Wilcoxon test (****$p < 0.0001$).

e, UMAP plot of indicated Tang et al.[23] scores in WCM1078 organoids treated with either 1μM A858 or 1μM A947 for 72h. Violin plots correspond to the UMAP plot above. Analyzed using the Wilcoxon test (****$p < 0.0001$).

f, UMAP plot of indicated Tang et al.[23] scores in MSK-PCa16 organoids treated with either 1μM A858 or 1μM A947 for 72h. Violin plots correspond to the UMAP plot above. Analyzed using Wilcoxon test (ns=not significant, *$p < 0.05$, **$p < 0.01$, ***$p < 0.001$, ****$p < 0.0001$).

g, Bubble plot of SCENIC results from WCM1078 organoids treated with either 1μM A858 or 1μM A947 for 72h.

h, Bubble plot of SCENIC results from MSK-PCa16 organoids treated with either 1μM A858 or 1μM A947 for 72h.

i, GSEA enrichment plots from WCM1078 of A947-treatment downregulated genes from scRNA-seq data.

j, GSEA enrichment plots from MSK-PCa16 of A947-treatment downregulated genes from scRNA-seq data.

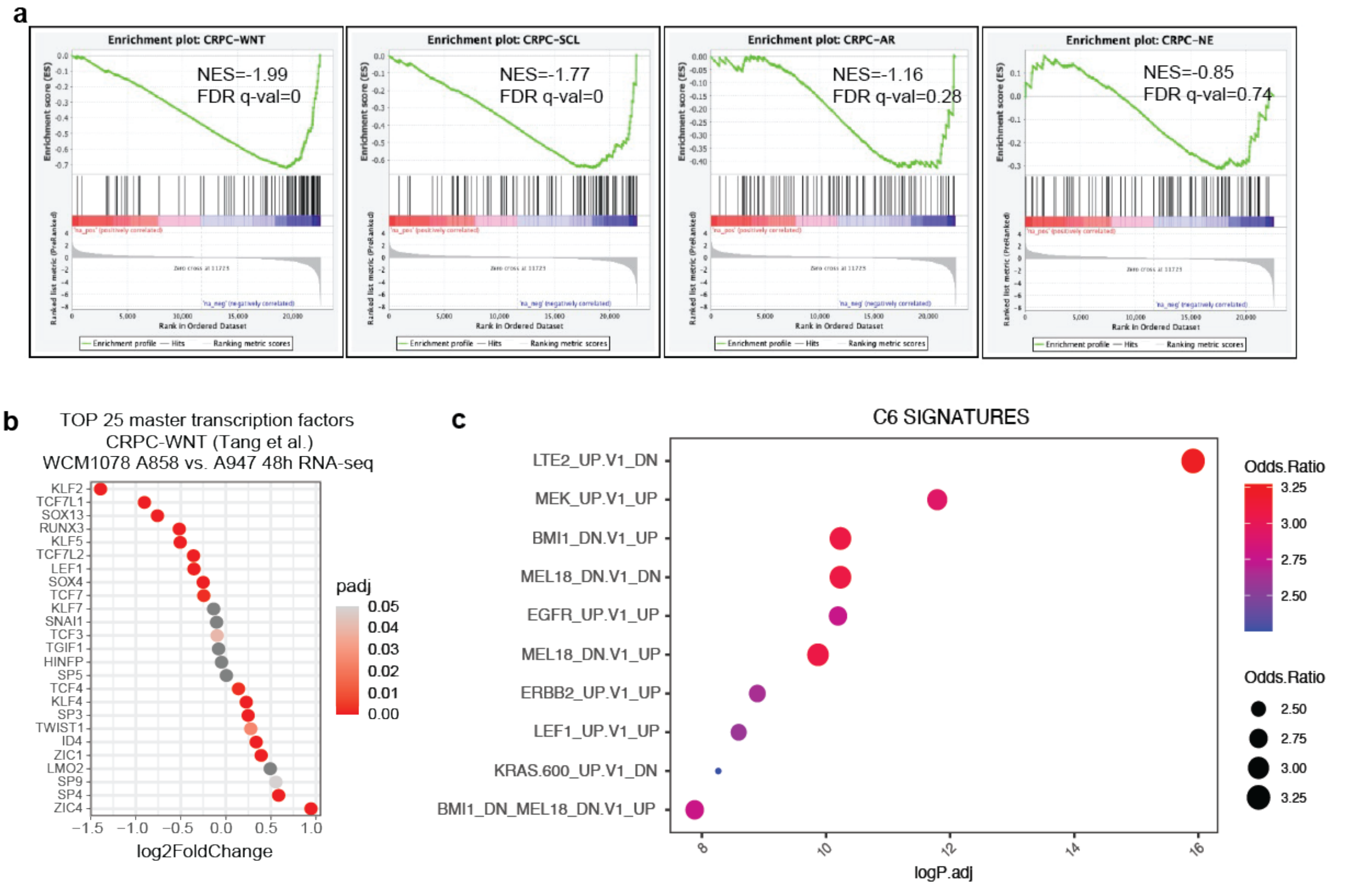


d

| ATAC-seq data | UP | DOWN | Unchanged | Total |
|---|---|---|---|---|
| LNCaP AU-15330 24h (Xiao et al.) | 1,104 | 2,242 | 45,264 | 48,810 |
| VCaP AU-15330 4h (Xiao et al.) | 3,396 | 5,347 | 41,910 | 55,716 |
| VCaP AU-15330 24h (Xiao et al.) | 11,079 | 10,833 | 33,804 | 50,653 |
| WCM1078 organoids A947 4h (this study) | 80 | 3,979 | 52,895 | 56,954 |

**Supplementary Figure 6. CRPC-WNT organoids become deregulated in multiple proliferative pathways upon SMARCA2/4 degradation**

a, GSEA enrichment plots from WCM1078 treated with 1μM A947 for 48h RNA-seq data.

b, Dot plot of expression levels of top 25 most active CRPC-WNT master transcription (determined by Tang et al[23]) from 1μM A947-treated WCM1078 (48h) bulk RNA-seq data.

c, Bubble plots from C6 signatures in WCM1078 treated with 1μM A947 for 48h RNA-seq data.

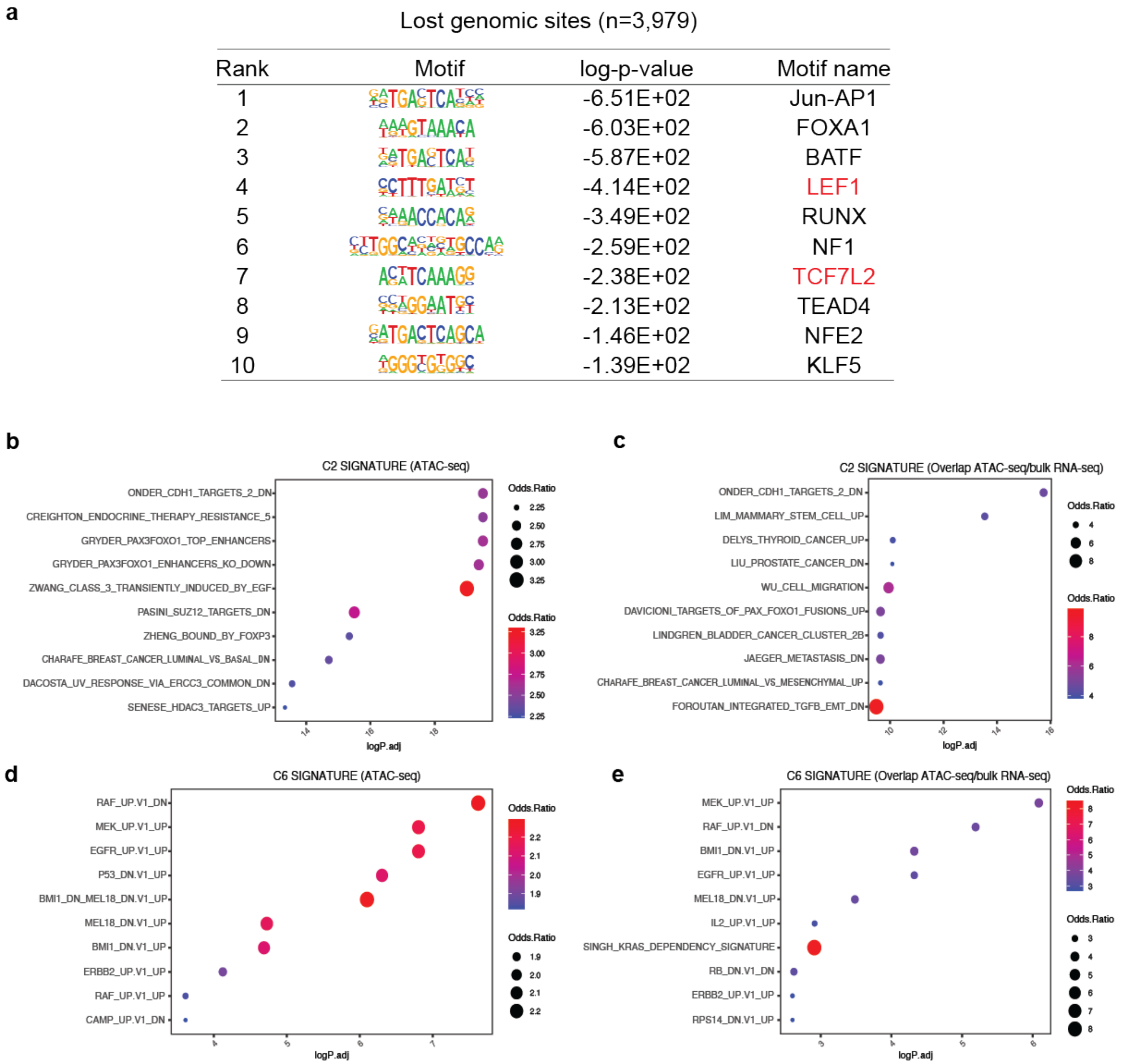


**Supplementary Figure 7. A947-treatment leads to decreased chromatin accessibility in CRPC-WNT**

a, Homer motif analysis from A947-treatment (4h) downregulated peaks in ATAC-seq data from WCM1078 organoids.

b, Bubble plots from C2 signatures in WCM1078 treated with 1μM A947 for 4h ATAC-seq data.

c, Bubble plots from overlap (ATAC-seq and RNA-seq 48h) of C2 signatures in WCM1078 treated with 1μM A947.

d, Bubble plots from C6 signatures in WCM1078 treated with 1μM A947 for 4h ATAC-seq data.

e, Bubble plots from overlap (ATAC-seq and RNA-seq 48h) of C6 signatures in WCM1078 treated with 1μM A947.

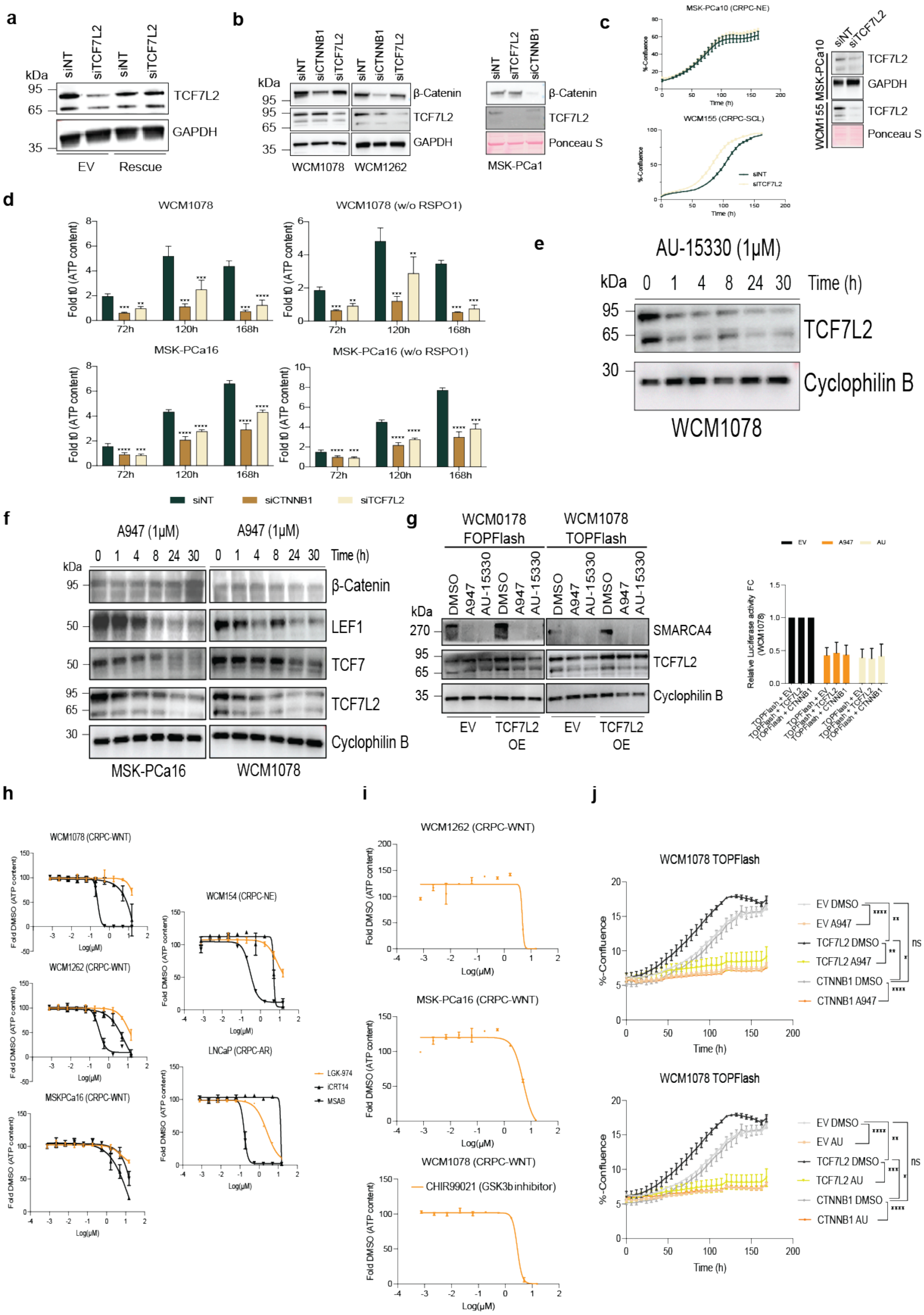


**Supplementary Figure 8. CRPC-WNT is dependent on TCF7L2**

a, Immunoblot of indicated proteins at 72h after transfection of indicated siRNA or plasmid (EV or rescue. GAPDH serves as loading control. Data is representative of *n* = 2 independent experiments. EV: empty vector.

b, Immunoblot of indicated proteins 72h after transfection with indicated siRNA. GAPDH or Ponceau S serve as loading controls. Data is representative of *n* = 2 independent experiments.

c, Immunoblot of indicated proteins 72h after transfection with indicated siRNA. GAPDH or Ponceau S serve as loading controls. Growth data measured by live-cell imaging (Incucyte S3). Data is representative of *n* = 2 independent experiments and are presented as mean values +/− SEM.

d, Proliferation was measured with CellTiter Glo 2.0 upon transfection of the indicated siRNA. Data are presented as mean values +/− SEM and analyzed using paired Students t-test (*p < 0.05, **p < 0.01, ***p < 0.001, ****p < 0.0001). Data is representative of *n* = 3 independent experiments.

e, Immunoblot of indicated proteins from WCM1078 organoid models upon treatment with AU-15330 over time. Cyclophilin B serves as loading control.

f, Immunoblot of indicated proteins from indicated organoids models upon treatment with A947 over time. Cyclophilin B serves as loading control.

g, Right: Immunoblot of indicated proteins from indicated in FOP/TOPFlash-transduced organoid models upon treatment with 1µM A947 and 1µM AU-15330 for 4h. Cyclophilin B serves as loading control. Left: Fold change (FC) of each TOPFlash condition shown in Figure 3d in relation to their OE condition. Data are presented as mean values +/− SEM.

h, Proliferation was measured with CellTiter Glo 2.0 upon treatment with indicated drugs. Data are presented as mean values +/− SEM). Data is representative of *n* = 2 independent experiments.

i, Proliferation was measured with CellTiter Glo 2.0 upon treatment with CHIR-99021. Data are presented as mean values +/− SEM.

j, Confluence measured with Incucyte S3 upon treatment with DMSO, 1µM A947 or 1µM AU-15330 (AU) in WCM1078 TOPFlash overexpressing empty vector (EV), TCF7L2 or CTNNB1. Data is representative of $n = 2$ independent experiments. Data are presented as mean values +/− SEM and analyzed using two-way ANOVA (*$p < 0.05$, **$p < 0.01$, ***$p < 0.001$, ****$p < 0.0001$).

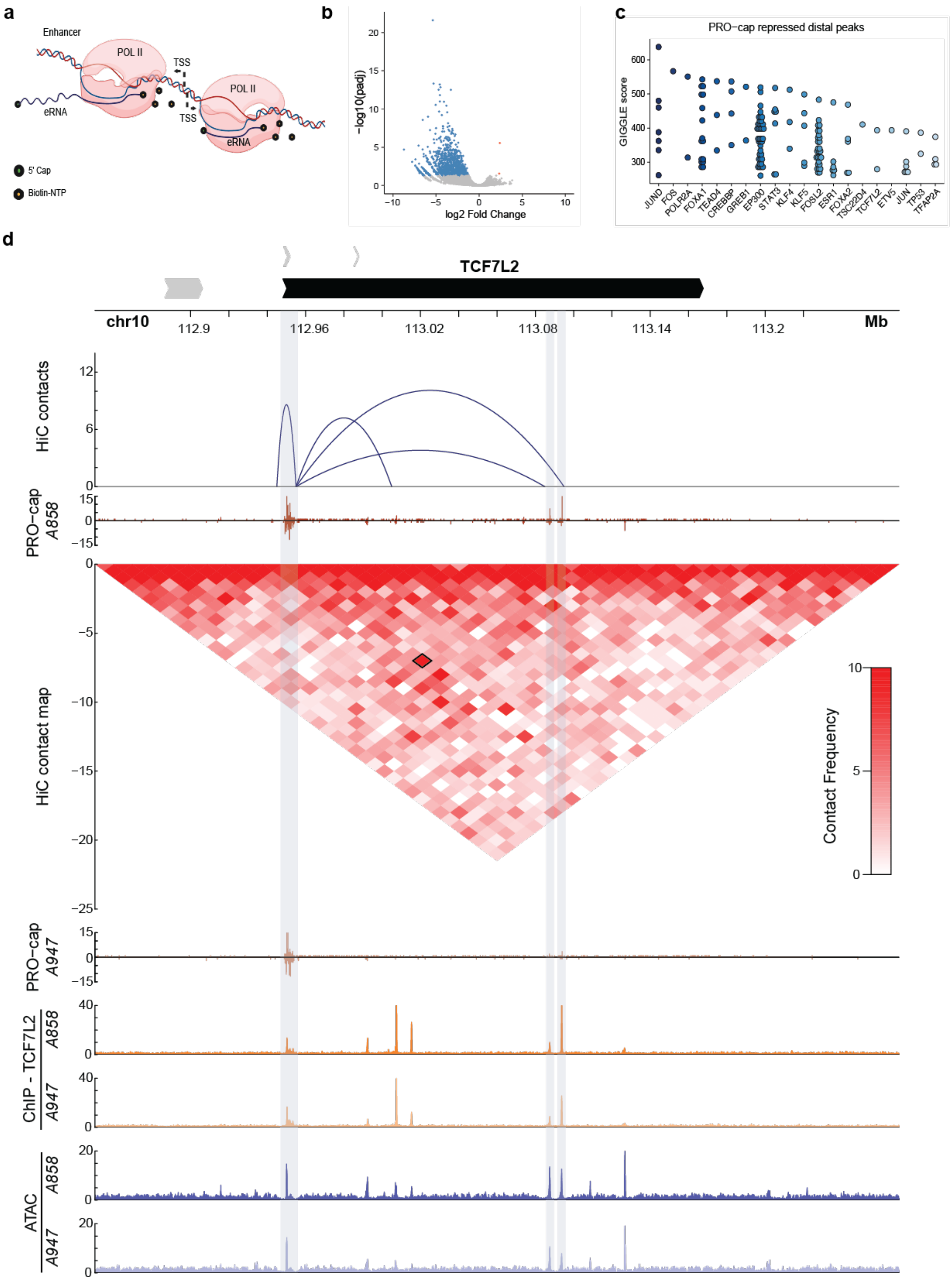

a
Enhancer
POL II
TSS
eRNA
5' Cap
Biotin-NTP
b
−log10(padj)
log2 Fold Change
c
PRO−cap repressed distal peaks
GIGGLE score
JUND
FOS
POLR2A
FOXA1
TEAD4
CREBBP
GREB1
EP300
STAT3
KLF4
KLF5
FOSL2
ESR1
FOXA2
TSC22D4
TCF7L2
ETV5
JUN
TP53
TFAP2A
d
TCF7L2
chr10
Mb
HiC contacts
PRO−cap A858
HiC contact map
Contact Frequency
PRO−cap A947
ChIP - TCF7L2 A858
A947
ATAC A858
A947

**Supplementary Figure 9. The TCF7L2 promoter interacts with an intragenic enhancer that is kept accessible by the SWI/SNF complex**

a, Model of PRO-cap: nascent RNA, like eRNA, is captured A nuclear run-on reaction with biotin-NTP and sarkosyl is carried out on nuclear lysates.

b, Volcano plot of differential expressed PRO-cap peaks A858 vs A947.

c, GIGGLE score from depleted distal PRO-cap peaks.

d, Hi-C heat maps from CRPC-WNT patient (from the Zhao et al. data set[44]**)** within the *TCF7L2* gene locus. PRO-cap, TCF7L2 ChIP-seq and ATAC-seq read-density tracks from the same A858 and A947 treatment conditions are overlaid. Grey highlights mark enhancers and the *TCF7L2* promoter. Loops indicate read-supported *cis* interactions within the locus.

**a**

Lost genomic sites (n=4,393)

| Rank | Motif | log-p-value | Motif name |
|---|---|---|---|
| 1 | 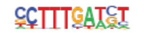 | -1.84E+03 | LEF1 |
| 2 | 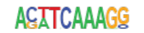 | -9.57E+02 | TCF7L2 |
| 3 | 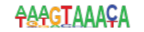 | -7.91E+02 | FOXA1 |
| 4 | 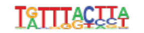 | -7.76E+02 | FOXM1 |
| 5 | 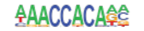 | -5.24E+02 | RUNX1 |
| 6 | 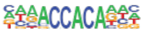 | -4.42E+02 | RUNX2 |
| 7 | 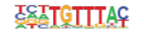 | -4.07E+02 | FOXK1 |
| 8 | 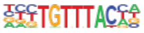 | -3.47E+02 | FOXP1 |
| 9 | 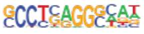 | -3.30E+02 | AP-2 |
| 10 | 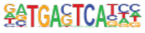 | -2.97E+02 | Jun-AP1 |

**b**

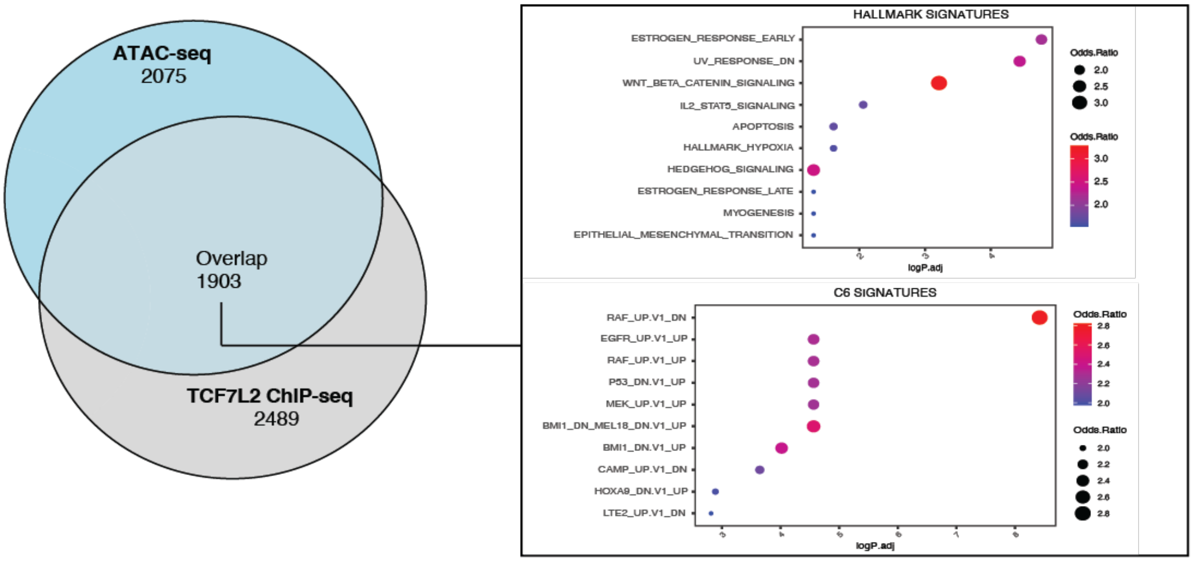


203 **Supplementary Figure 10. TCF7L2 chromatin binding is reduced upon treatment with A947 in CRPC-WNT**

a, Homer motif analysis from A947-treatment (4h) downregulated peaks in TCF7L2 ChIP-seq data from WCM1078 organoids.

b, Left, Venn diagram depicting overlap between ATAC-seq and TCF7L2 ChIP-seq data A947-treatment downregulated peaks. Right, GSEA depicting HALLMARK and C6 signatures from overlap of ATAC-seq and TCF7L2 ChIP-seq A947-treatment downregulated peaks.

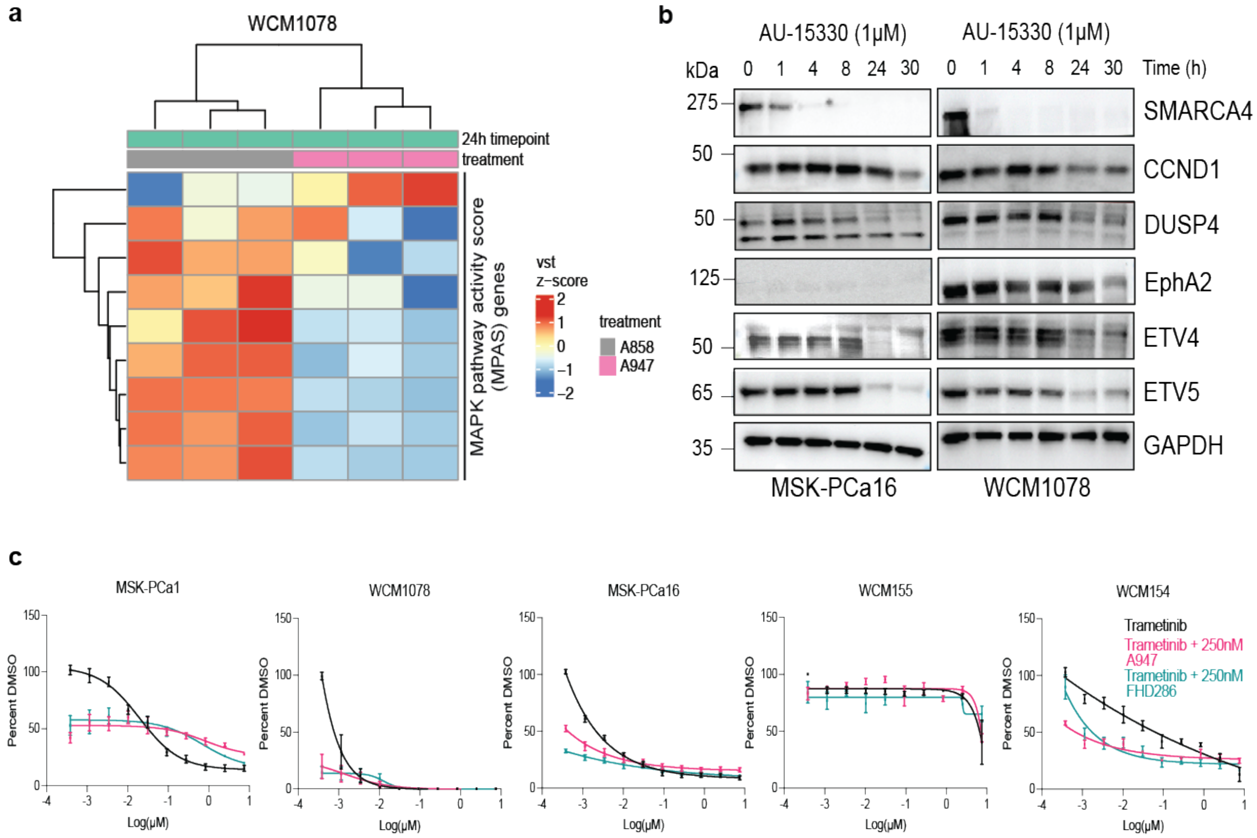


**Supplementary Figure 11. TCF7L2 regulates pro-proliferative pathways in CRPC-WNT**

a, RNA-seq heat maps for MPAS genes in WCM1078 with or without 24 h of A947 treatment.

b, Immunoblot of indicated proteins at indicated time upon treatment with 1μM AU-15330. GAPDH serves as loading control. Data is representative of *n* = 2 independent experiments.

c, Celltiter Glo 2.0 assay growth curves measure after 7 days of treatment with indicated drugs. Data are presented as mean values +/− SEM of *n* = 3 independent experiments.

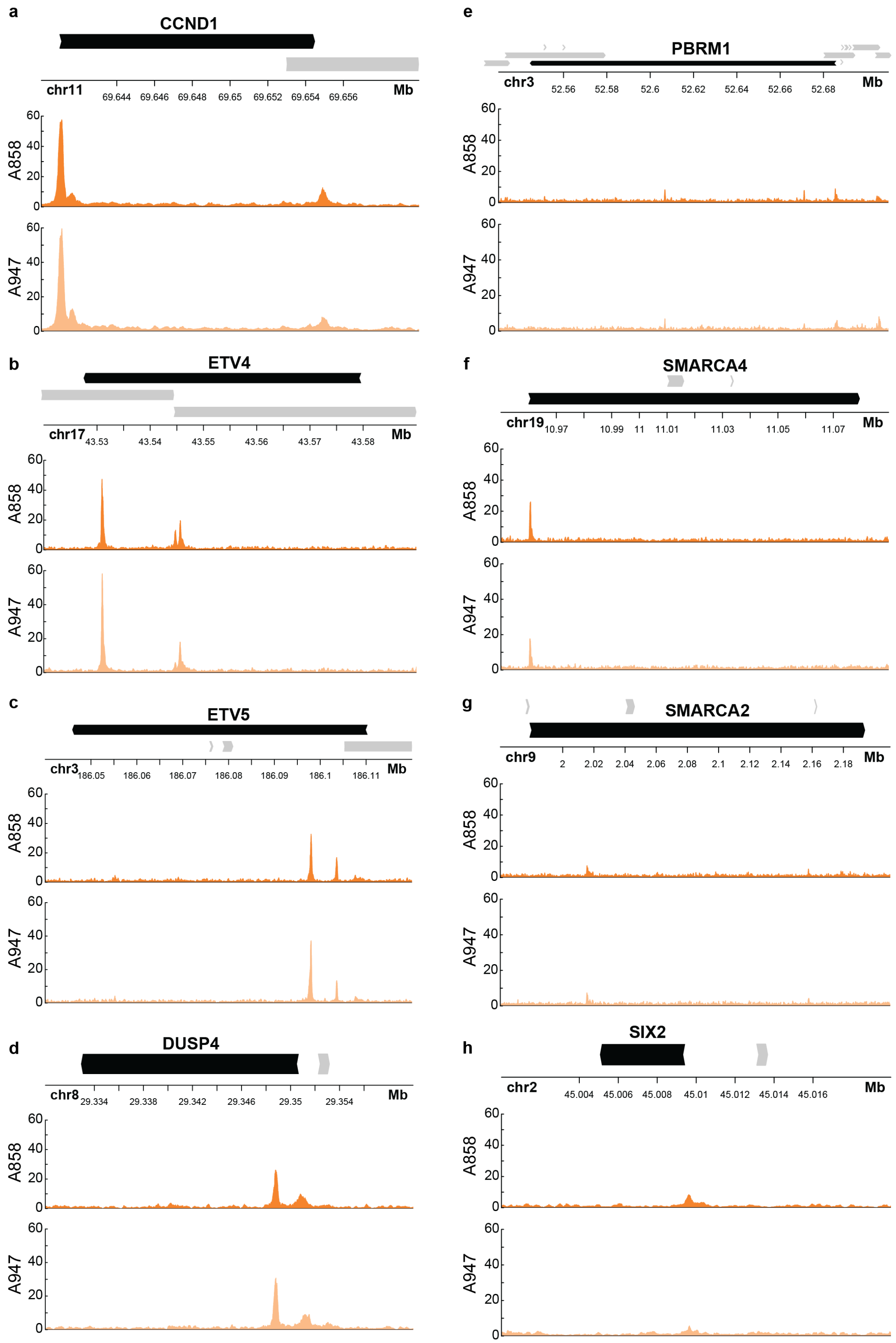
a
CCND1
chr11
69.644 69.646 69.648 69.65 69.652 69.654 69.656
Mb
A858
A947
0 20 40 60
b
ETV4
chr17
43.53 43.54 43.55 43.56 43.57 43.58
Mb
A858
A947
c
ETV5
chr3
186.05 186.06 186.07 186.08 186.09 186.1 186.11
Mb
A858
A947
d
DUSP4
chr8
29.334 29.338 29.342 29.346 29.35 29.354
Mb
A858
A947
e
PBRM1
chr3
52.56 52.58 52.6 52.62 52.64 52.66 52.68
Mb
A858
A947
f
SMARCA4
chr19
10.97 10.99 11 11.01 11.03 11.05 11.07
Mb
A858
A947
g
SMARCA2
chr9
2 2.02 2.04 2.06 2.08 2.1 2.12 2.14 2.16 2.18
Mb
A858
A947
h
SIX2
chr2
45.004 45.006 45.008 45.01 45.012 45.014 45.016
Mb
A858
A947

**Supplementary Figure 12. TCF7L2 chromatin interactions**

a, UCSC browser tracks at *CCND1* gene loci and TCF7L2 ChIP-seq tracks.

b, UCSC browser tracks at *ETV4* gene loci and TCF7L2 ChIP-seq tracks.

c, UCSC browser tracks at *ETV5* gene loci and TCF7L2 ChIP-seq tracks.

d, UCSC browser tracks at *DUSP4* gene loci and TCF7L2 ChIP-seq tracks.

e, UCSC browser tracks at *PBRM1* gene loci and TCF7L2 ChIP-seq tracks.

f, UCSC browser tracks at *SMARCA4* gene loci and TCF7L2 ChIP-seq tracks.

g, UCSC browser tracks at *SMARCA2* gene loci and TCF7L2 ChIP-seq tracks.

h, UCSC browser tracks at *SIX2* gene loci and TCF7L2 ChIP-seq tracks.
